# A bacterial Argonaute with efficient DNA and RNA cleavage activity guided by small DNA and RNA

**DOI:** 10.1101/2021.10.11.464003

**Authors:** Longyu Wang, Xiaochen Xie, Yang Liu, Wenqiang Li, Bin Lv, Zhiwei Zhang, Jun Yang, Guangbo Yan, Wanping Chen, Cheng Zhang, Fei Wang, Lixin Ma

**Affiliations:** State Key Laboratory of Biocatalysis and Enzyme Engineering, Hubei Collaborative Innovation Center for Green Transformation of Bio-resources, Hubei Key Laboratory of Industrial Biotechnology, School of Life Sciences, Hubei University, Wuhan, Hubei 430062, China

## Abstract

Argonaute proteins are widespread in prokaryotes and eukaryotes. Most prokaryotic Argonaute proteins (pAgos) use 5’P-gDNA to target complementary DNA. However, more and more studies on the properties of pAgos make their functions more diversified. Previously reported pAgos only possess several forms of high activity in all eight cleavage patterns, which limits their practical applications. Here, we described a unique pAgo from *Marinitoga hydrogenitolerans* (MhAgo) with eight cleavage activities. MhAgo can utilize all four types of guides (5’OH-gDNA, 5’P-gDNA, 5’OH-gRNA, and 5’P-gRNA) for ssDNA and RNA cleavage. Further studies demonstrated that MhAgo had high activities with 16-21 nt guides and no obvious preferences for the 5’-end nucleotides of 5’OH-guides. Unexpectedly, MhAgo had different preferences for the 5’-end nucleotides of 5’P-guides depending on the types of targets. Although the specificity of MhAgo was related to the types of guides, single mismatches in the central and 3’-supplementary regions of guides greatly reduced the cleavage efficiency. Additionally, the electrophoretic mobility shift assay (EMSA) demonstrated MhAgo had the weakest affinity for 5’P-gRNA:tRNA duplex, which was consistent with its cleavage efficiency. In conclusion, MhAgo is highly active under a wide range of conditions and can be used for programmable endonucleolytic cleavage of both ssDNA and RNA substrates. The abundant biochemical characteristics of MhAgo broaden our understanding of pAgos and expand the potential application in nucleic acids manipulations.

## INTRODUCTION

Argonaute proteins (Agos) are distributed in eukaryotes, bacteria and archaea (1). In eukaryotes, Agos act in a complex with small non-coding RNAs to implement RNA interference (RNAi) (2). Eukaryotic Argonaute proteins (eAgos) not only participate in the biogenesis of small regulatory RNAs, but also regulate gene expression and defend against foreign pathogen invasion via RNAi (1,3). In general, eAgos use 5’ phosphorylated RNA guides (5’P-gRNA) to target complementary RNA (4). However, due to the lack of RNAi pathway, the function of prokaryotic Argonaute proteins (pAgos) in bacteria and archaea remains poorly understood. Encouragingly, recent studies show that pAgos can function in host defense against invasive genetic elements like plasmids and phages and may be involved in a variety of activities within the cell, like DNA replication (5–9).

Structural studies have revealed that some Agos adopt the four-domain architecture, including N-terminal (N), PIWI-Argonaute-Zwille (PAZ), middle (MID) and catalytic RNase H-like P element-induced wimpy testis (PIWI) domains. The PAZ domain anchors the 3’ end of the nucleic acid guide by bending it into a specific binding pocket. The MID domain anchors the 5’ end of the nucleic acid guide by providing a binding pocket. The active center of Agos is in the PIWI domain, which is structurally similar to ribonuclease H (RNase H) (10–12). Agos with DEDX (X = N, D or H) catalytic tetrad in PIWI domains generally possess endonuclease activity (13). Despite the structural homology between pAgos and eAgos, most pAgos utilize DNA guides (gDNA) to target complementary DNA in contrast to eAgos (14,15).

Most of the currently reported pAgos possess small DNA guided ssDNA cleavage activity, including TtAgo (*Thermus thermophilus*) (15), PfAgo (*Pyrococcus furiosus*) (16), MjAgo (*Methanocaldococcus jannaschii*) (17,18), CbAgo (*Clostridium butyricum*) (19,20), LrAgo (*Limnothrix rosea*) (20), CpAgo (*Clostridium perfringens*) (21), IbAgo (*Intestinibacter bartlettii*) (21), SeAgo (*Synechococcus elongatus*) (8), CbcAgo (*Clostridium butyricum CWBI1009*) (22), KmAgo (*Kurthia massiliensis*) (23,24) and FpAgo (Ferroglobus placidus) (25). All these pAgos have 5’ phosphorylated DNA guides (5’P-gDNA) mediated ssDNA cleavage activity, but only CbAgo (19,20), LrAgo (20), CpAgo (21), IbAgo (21), SeAgo (8) and KmAgo (23,24) are reported to have 5’ hydroxylated DNA guides (5’OH-gDNA) mediated ssDNA cleavage activity. Some pAgos can use small RNA guides (gRNA) to target nucleic acids, including AaAgo (*Aquifex aeolicus*) (26), MpAgo (*Marinitoga piezophila*) (27), TpAgo (*Thermotoga profunda*) (27) and KmAgo (23,24). Additionally, some pAgos can use RNA as substrate, including AaAgo (26), TtAgo (15), MpAgo (27), CpAgo (21) and KmAgo (23,24). Not all pAgos have cleavage activity, some pAgos only have binding activity but no cleavage activity, such as RsAgo (*Rhodobacter sphaeroides*) (28) and AfAgo (*Archaeoglobus fulgidus*) (29). Different from many pAgos, RsAgo specifically recognizes 5’P-RNA as the guide strand and DNA as the target strand (28) (Supplementary Table S1). The nucleic acid-guided cleavage activities of Argonaute proteins are reminiscent of the Clustered Regularly Interspaced Short Palindromic Repeats (CRISPR) - associated proteins (Cas) (30,31). Like CRISPR-associated proteins, pAgos may potentially be used in nucleic acid detection (32–36), molecular cloning (37) and genome editing application (38). The pAgos may have the following activities: 5’OH-gDNA or 5’P-gDNA mediated ssDNA cleavage activity, 5’ hydroxylated RNA guides (5’OH-gRNA) or 5’P-gRNA mediated ssDNA cleavage activity, 5’OH-gDNA or 5’P-gDNA mediated RNA cleavage activity and 5’OH-gRNA or 5’P-gRNA mediated RNA cleavage activity. The most omnipotent Ago protein that has been reported is KmAgo which has 5’OH-gDNA or 5’P-gDNA mediated cleavage activity, and relatively weak 5’P-gRNA mediated cleavage activity. Moreover, KmAgo is unobserved with 5’OH-gRNA mediated cleavage activity (23,24).

In 2016, Emine Kaya et al found a new class of pAgos which were associated with CRISPR–Cas containing MpAgo and TpAgo (27,39,40). They preferentially utilize 5’OH-gRNA that are chemically distinct from the 5’P-gDNA used by other pAgos to target nucleic acids. In order to discover more properties of pAgos, we chose the protein sequence of MpAgo as the query to search for pAgos. Finally, we selected the pAgo from *Marinitoga hydrogenitolerans* (MhAgo) to study. Like MpAgo, MhAgo prefers to bind 5’OH-gRNA for complementary target cleavage. Unexpectedly, MhAgo has eight patterns of cleavage activity including 5’OH-gDNA or 5’P-gDNA mediated ssDNA/RNA cleavage activity, 5’OH-gRNA or 5’P-gRNA mediated ssDNA/RNA cleavage activity. Furthermore, MhAgo can cleave DNA target with high efficiency mediated by 5’P-gDNA, 5’OH-gRNA and 5’P-gRNA, and RNA target with high efficiency mediated by 5’OH-gDNA, 5’P-gDNA and 5’OH-gRNA. Moreover, in comparison with most reported pAgos which have no obvious cleavage preference for the 5’-terminal nucleotides of guides, the cleavage preference for the 5’-terminal nucleotides of guides of MhAgo is affected by the nature of guides and targets. Meanwhile, MhAgo has good specificity for single mismatches that occurred in the central and 3’-supplementary regions (position 9-15). The abundant biochemical characteristics of MhAgo broaden our understanding of pAgos and make it have a wider application prospect.

## MATERIAL AND METHODS

### Protein expression and purification

The MhAgo gene was synthesized by Wuhan Genecreate, was codon-optimized for expression in *E. coli* and inserted into a pET28a expression vector in frame with the N-terminal His tag. The MhAgo protein was expressed in *E. coli* strain BL21(DE3). For protein expression, cells were grown in LB medium containing 50 μg/ml kanamycin to an OD_600_ of 0.8, expression was induced by the addition of IPTG (isopropyl β-D-1-thiogalactopyranoside) to 0.5 mM final concentration, and cells were incubated at 18 °C while shaking for 20 h. Cells were harvested by centrifugation and resuspended in buffer A (20 mM HEPES, pH 7.5, 250 mM NaCl, 1 mM DTT), and supplemented with an ethylenediaminetetraacetic acid (EDTA)-free protease inhibitor cocktail tablet (Roche). Cells were lysed via high-pressure homogenization, and the lysate was clarified by centrifugation at 18 000 rpm for 50 min. Then the supernatant was bound to Ni-NTA agarose resin. The column was washed with buffer A containing 20-100 mM imidazole with 30 column volumes, and the proteins were eluted by increasing gradually the concentration of imidazole in buffer A to 1 M. The eluted protein was concentrated by ultrafiltration using an Amicon 50K filter unit (Millipore). Next, the concentrated protein was diluted with 20 mM HEPES, pH 7.5 to lower the final salt concentration to 50 mM NaCl. The diluted protein was applied to a 5-mL Heparin HiTrap column (GE Life Sciences), washed with buffer B (20 mM HEPES, pH 7.5, 50 mM NaCl) and eluted with a linear gradient of 0.125–2 M NaCl. Then the heparin-purified protein was concentrated by ultrafiltration and applied to a size exclusion column (Superdex 200 16/600 column, GE Healthcare). The protein was eluted with Buffer C (20 mM HEPES, pH 7.5, 250 mM NaCl). Eluted protein was concentrated, flash-frozen in liquid nitrogen, and stored at −80 °C.

### The activity assays of ssDNA and RNA cleavage

The nucleic acid guides and targets including 5’-FAM-labeled targets and 5’-P-labeled guides were synthesized by Sangon and Genscript (Supplementary Table S2). Unless otherwise indicated, purified MhAgo, ssDNA or RNA guides, and ssDNA or RNA targets were generally mixed in 4:2:1 ratio (800 nM MhAgo: 400 nM guide: 200 nM target) in 5× Reaction Buffer (50 mM HEPES–NaOH pH 7.5, 500 mM NaCl, 50 mM MnCl_2_, 50% glycerol). The targets were added after that MhAgo and guides had been incubated for 10 min at 55 °C. Then reaction mixtures were incubated for 30 min at 60 °C. The lengths and 5’-end nucleotides of guides were pointed out in the text or legend. The incubation temperature, incubation time and divalent metal ion concentration were varied in different experiments. To analyze the effect of various divalent cations on cleavage activity, 5 mM Mg^2+^, Ni^2+^, Co^2+^, Cu^2+^, Fe^2+^, Ca^2+^ or Zn^2+^ were added instead of Mn^2+^ and reaction for 30 min at 55 °C. When studying the effect of mismatches on cleavage activity, the incubation time of 5’OH-gDNA mediated ssDNA cleavage assays was 1 h. All reactions were terminated by adding an equal volume of RNA loading dye (95% formamide, 18 mM EDTA, 0.025% SDS and 0.025% bromophenol blue). Then the samples were incubated for 5 min at 95 °C before resolving on 20% denaturing polyacrylamide gels. Nucleic acids were stained using SYBR gold Nucleic Acid Gel Stain (Invitrogen) and visualized using Gel Doc^™^ XR+ (Bio-Rad). Cleavage experiments were tested in three independent experiments. The percentage of cleavage was analyzed using ImageJ and Prism 8 (GraphPad) softwares. The ssDNA and RNA targets used in quantitative experiments were FAM-labeled and the others were non-labeled.

### Electrophoretic mobility shift assay (EMSA)

To analyze the affinity of MhAgo to different guides, MhAgo and different types of guides were incubated in 10 μl of binding buffer containing 10 mM HEPES–NaOH pH 7.5, 100 mM NaCl, 5 mM MnCl_2_, 5% glycerol for 30 min at 55 °C. The concentration of guides was fixed as 128 nM, whereas the concentration of MhAgo varied. After incubation, the samples were mixed with 10× non-denaturing loading buffer containing 250 mM Tris-HCl (pH 7.5), 40% glycerol and 0.1% bromophenol blue. The mixed samples were resolved by 10% native PAGE with 0.5× Tris-Borate-EDTA (TBE) buffer for 1 h at 120 V. The free nucleic acids and MhAgo/guides complexes were visualized using Gel Doc^™^ XR+.

To analyze the affinity of MhAgo to different guide-target duplexes, guides and targets were firstly annealed to form duplexes. The reactions were performed at 65 °C for 3 min, decreased to 25 °C at a rate of 0.1 °C/s and maintained for 2 min. Then MhAgo_DM and different guide-target duplexes were incubated in 10 μl of binding buffer containing 10 mM HEPES–NaOH pH 7.5, 100 mM NaCl, 5 mM MnCl_2_, 5% glycerol for 30 min at 55 °C. The concentration of nucleic acid duplexes was fixed as 64 nM, whereas the concentration of MhAgo varied. After incubation, the samples were mixed with 10× non-denaturing loading buffer and resolved by 10% native PAGE. Free nucleic acids and MhAgo/duplexes complexes were visualized using Gel Doc^™^ XR+.

Binding experiments were tested in three independent experiments. The percentage of binding to guides and duplexes was analyzed using ImageJ and Prism 8 (GraphPad) software. The data were fitted with the Hill equation with a Hill coefficient of 2–2.5.

### Co-purification nucleic acids

Proteinase K (Ambion) of final concentrations of 1 mg/mL was added to 2 mg of Ni-NTA purified MhAgo. The sample was incubated for 1 h at 55 °C. The nucleic acids were separated from the organic fraction by adding Roti-phenol/chloroform/isoamyl alcohol pH 7.5-8.0 in a 1:1 ratio and centrifugation at 12,000 rpm for 15 min. The top layer was isolated and nucleic acids were precipitated by adding 99% ethanol in a 1:2 ratio. This mixture was incubated for 4 h at −20 °C and centrifuged at 12,000 rpm for 30 min. Next, the nucleic acid pellet was washed with 700 μl of 70% ethanol and solved in 20 μl nuclease-free water. The purified nucleic acids were treated with either 100 μg/ml RNase A (Thermo Fisher Scientific), 2 units DNase I (NEB), or both for 30 min at 37 °C. Finally, the samples were resolved on a denaturing urea polyacrylamide gel (20%) and stained with SYBR Gold. The co-purified nucleic acids were visualized using Gel Doc^™^ XR+.

### Mutant construction and structure analysis of MhAgo

The mutants of MhAgo including MhAgo_DM, mutant_I386V/A, mutant_F378Y/A, mutant_L379I/A and mutant_I386V/F378Y/L379I were constructed by PCR-mediated site-directed mutagenesis (41). The mutants were expressed and purified like wild-type MhAgo. The homology model of MhAgo was performed with SWISS-MODEL using the binary structure of MpAgo and 5’OH-gRNA (PDB ID: 5I4A, identity 65.41%) as the template. The structure alignments of MhAgo and other Agos were analyzed using PyMol software.

## RESULTS

### MhAgo exhibits eight patterns of cleavage activity

MpAgo preferentially uses 5’OH-gRNA to cleave complementary ssDNA and RNA target (27). In order to discover more properties of pAgos, we chose the protein sequence of MpAgo as the query and used the web interface of the BLASTp program to search for new pAgos. Finally, we selected one pAgo originated from *Marinitoga hydrogenitolerans* (MhAgo) which shared 65.41% sequence identity with MpAgo. Phylogenetic analysis revealed that MhAgo indeed was closely related to MpAgo (Figure 1A). The target cleavage activity of Agos is related to the conserved DEDX tetrad in the PIWI domain (10). Detailed sequence alignments showed that both MhAgo and MpAgo had conserved catalytic tetrad (DEDN) in the PIWI domain (Supplementary Figure S1A).

**Figure 1.**
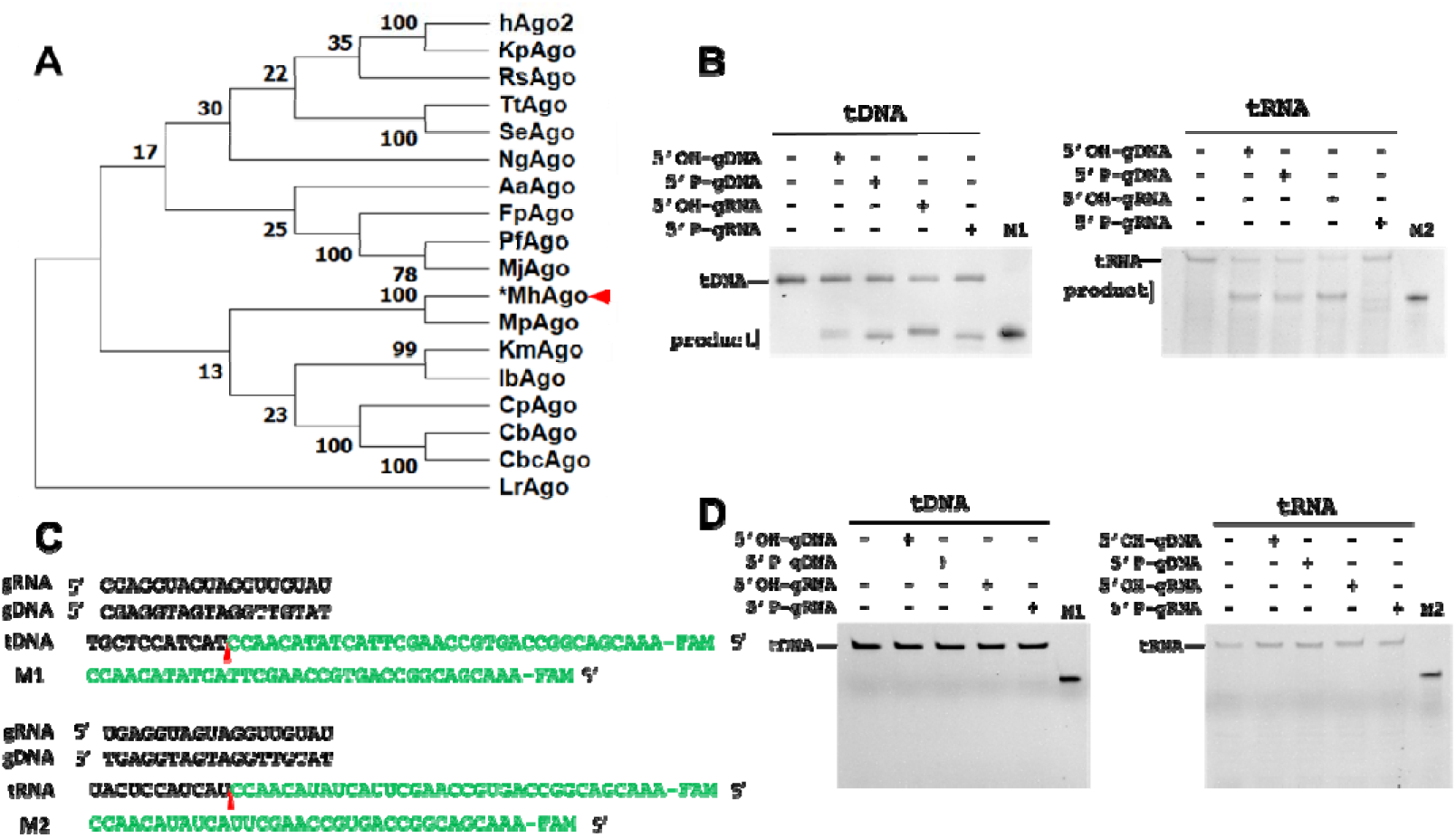
MhAgo exhibits eight patterns of cleavage activity. (A) Maximum likelihood phylogenetic tree of MhAgo and other previously characterized Agos. (B) Cleavage activity assay with FAM-labeled DNA and RNA targets using different guides. (C) The sequence of synthetic guides and targets used in cleavage activity assay. Cleavage product sequences are marked in green. The red triangle indicates the predicted cleavage site. (D) Substitutions of two out of four catalytic tetrad residues (D527A/D596A) causes the loss of MhAgo activity. MhAgo or MhAgo_DM, guides and targets were mixed in a 4:2:1 molar ratio (800 nM MhAgo or MhAgo_DM preloaded with 400 nM guide, plus 200 nM target) and incubated for 30 min at 60 °C, and the reaction buffer contained 5 mM Mn^2+^. M1 indicates synthesized 5’ FAM-labeled, 34-nt ssDNA product. M2 indicates synthesized 5’ FAM-labeled, 34-nt RNA product. The ssDNA targets are completely complementary to corresponding guides the 5’-end nucleotides of which are C. The RNA targets are completely complementary to corresponding guides the 5’-end nucleotides of which are T or U.

To study the biochemical properties of MhAgo, we purified MhAgo (Supplementary Figure S1B) and performed *in vitro* cleavage assay. Firstly, MhAgo was loaded with synthetic either DNA or RNA guides (18 nucleotides in length) including 5’P or 5’OH-oligonucleotides for 10 min at 55 °C. Then synthetic fluorescently labeled ssDNA or RNA targets were added and the mixture was incubated for 30 min at 60 °C in reaction buffer containing 5 mM Mn^2+^. The 20% denaturing gel showed that substrate cleavage by MhAgo was observed in all eight combinations of guide and target molecules (5’OH-gDNA-tDNA, 5’P-gDNA-tDNA, 5’OH-gRNA-tDNA, 5’P-gRNA-tDNA, 5’OH-gDNA-tRNA, 5’P-gDNA-tRNA, 5’OH-gRNA-tRNA, 5’P-gRNA-tRNA). We did not observe cleavage of the ssDNA or RNA substrates in the absence of guides (Figure 1B). The RNA cleavage guided by 5’P-RNA couldn’t be obviously observed, but obvious cleavage can be observed in mismatches experiments (Supplementary Figure S5D). The sequences of targets and guides used in the activity assay were shown in Figure 1C. When using 5’OH-gDNA or 5’OH-gRNA to target ssDNA, the cleavage site of the target strand occurred between 10th and 11th nucleotides counting from the 5’ end of guides, resulting in the appearance of the 34 nt long 5’-fragment of target DNA. When guided by 5’P-DNA, the cleavage site of the target strand occurred between 11th and 12th nucleotides counting from the 5’ end of guides, resulting in the appearance of the 33 nt long 5’-fragment of target DNA. When guided by 5’P-RNA, target cleavage occurred at 1–2 nucleotides downstream of the canonical cleavage site, resulting in the appearance of the 33 nt long and a few shorter 5’-fragment of target DNA (Supplementary Figure S1C, upper panel). However, when targeting RNA with 5’OH-gDNA, 5’P-gDNA and 5’OH-gRNA, the cleavage sites of the target strands were identical and occurred between 10th and 11th nucleotides counting from the 5’ end of guides, resulting in the appearance of the 34 nt long 5’-fragment of target RNA. When guided by 5’P-RNA, target cleavage occurred at 1-2 nucleotides downstream of the canonical cleavage site, resulting in the appearance of the 33 nt long and shorter 5’-fragment of target RNA (Supplementary Figure S1C, lower panel). The cleavage positions of MhAgo and MpAgo shifted from the canonical cleavage site when using 5’P-guides, which was also observed for some other Argonaute proteins using 5’OH-guides, including hAgo2 (Homo sapiens) (42), MjAgo (17), LrAgo (20), CpAgo and IbAgo (21).

The DEDX catalytic site in the PIWI domain is vital to the endonucleolytic activity of Argonaute proteins (27). In the case of MhAgo, this concerned residues D445, E481, D515 and N623. We obtained the catalytically inactive mutant of MhAgo (MhAgo_DM) with substitutions of two out of four catalytic tetrad residues with alanine (D445A/D515A). With regarding all eight combinations of guides and targets, no cleavage was detected for a catalytically dead MhAgo variant (Figure 1D). Furthermore, we analyzed the nucleic acids that co-purified with MhAgo expressed in *E. coli.* Denaturing polyacrylamide gel electrophoresis revealed that MhAgo co-purified nucleic acids were susceptible to DNase

I but not to RNase A treatment (Supplementary Figure S1D), indicating that MhAgo could acquire DNA fragments *in vivo.* But we don’t exclude that MhAgo may also target RNA *in vivo* which requires further investigation.

The eight cleavage activities of MhAgo have not been observed in other reported pAgo homologs. Distinct from the most reported pAgos, MhAgo can use both DNA and RNA guides for efficient cleavage. The biochemical properties of MhAgo need to be systematically characterized.

### Effects of guide lengths on target cleavage

Previously reported pAgos can function over a wide range of guide lengths, recently reported FpAgo has cleavage activity only guided by short DNA of 15–20 nt lengths (25). We next tested MhAgo cleavage efficiency using guides ranging from 12 to 40 nt lengths which shared identical sequences at their 5’ end. The optimal lengths of guides varied for different binding forms of guides and targets. For 5’OH-gDNA mediated DNA cleavage, 14-30 nt guides led to similar cleavage efficiency, while shorter guides rapidly diminished the cleavage efficiency (Figure 2A, upper panel). For 5’OH-gDNA mediated RNA cleavage, the cleavage was observed directed by 13-30 nt long guides and MhAgo was most active with 14-19 nt guides, with a lower cleavage efficiency observed with shorter or longer guides (Figure 2A, lower panel). For 5’OH-gRNA mediated target cleavage, the cleavage was observed directed by 13-30 nt long guides. For DNA target, the 5’OH-gRNA between 16 and 30 nt long resulted in similar cleavage activity (Figure 2B, upper panel). For RNA target, MhAgo could efficiently cleave targets only with 5’OH-gRNA between 16 and 21 nt (Figure 2B, lower panel). For 5’P-gDNA mediated target cleavage, the cleavage was observed directed by 13-40 nt long guides and the 5’P-gDNA between 18 and 30 nt long resulted in higher cleavage activity (Figure 2C). For 5’P-gRNA mediated DNA cleavage, the cleavage was observed directed by 14-30 long guides (Figure 2D, upper panel). For 5’P-gRNA mediated RNA cleavage, the cleavage was observed directed by 16-25 long guides (Figure 2D, lower panel). The 5’P-gRNA between 16 and 21 nt long resulted in higher cleavage activity for both DNA and RNA targets (Figure 2D). The precise target cleavage by MhAgo can be observed with 5’OH-gDNA, 5’OH-gRNA and 5’P-gDNA (Figure 2A-C). When using 5’P-gRNA to target DNA, the precise cleavage was observed with 14, 15 nt guides and the cleavage position shifted by one nucleotide with longer guides (Figure 2D, upper panel). When using 5’P-gRNA to target RNA, the precise cleavage was observed with 16, 17 nt guides and the cleavage position shifted by one nucleotide with longer guides (Figure 2D, lower panel).

**Figure 2.**
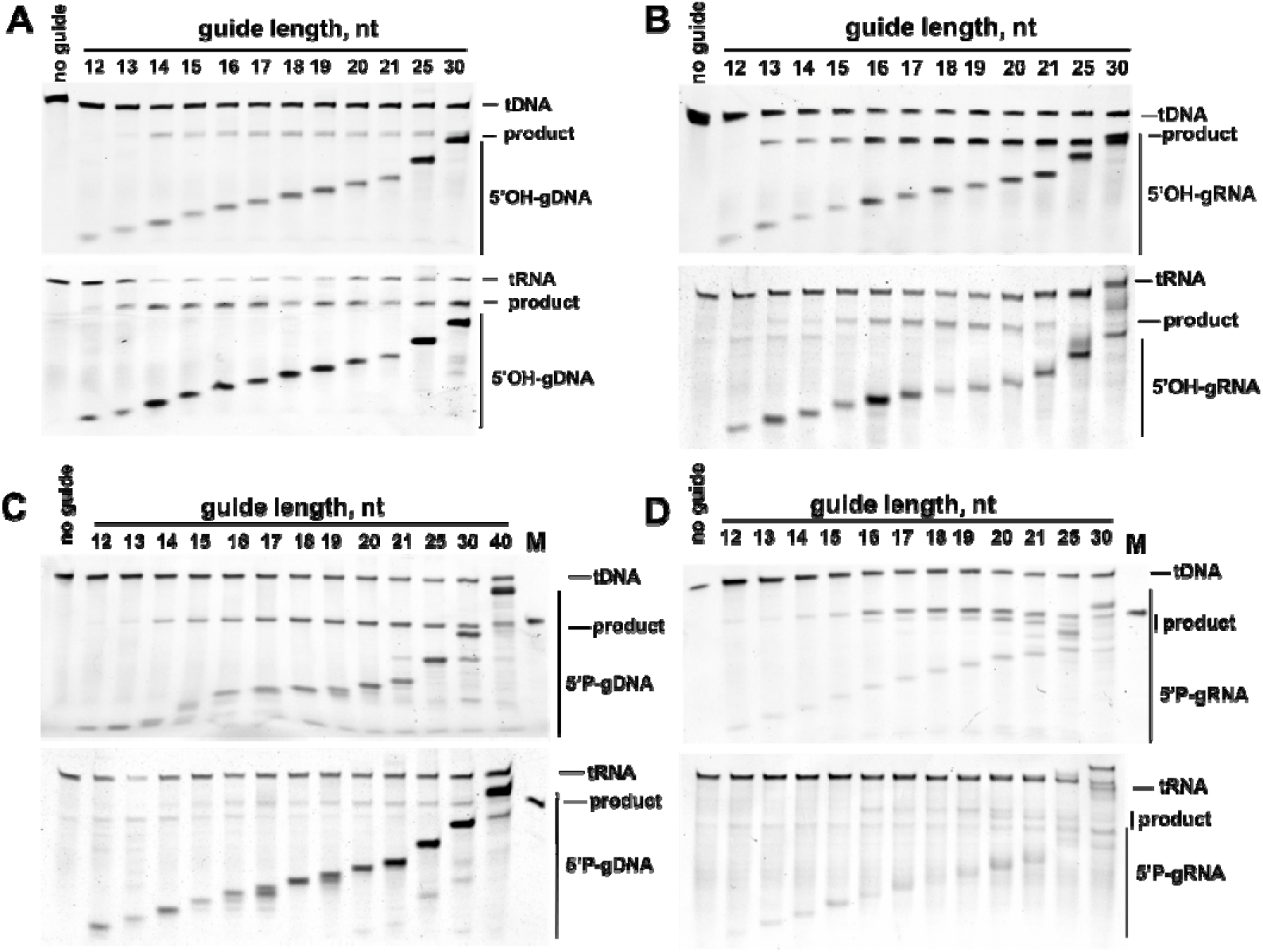
Effects of guide lengths on MhAgo activity. (A) Effects of the 5’OH-gDNA lengths on MhAgo cleavage activity. (B) Effects of the 5’OH-gRNA lengths on MhAgo cleavage activity. (C) Effects of the 5’P-gDNA lengths on MhAgo cleavage activity. (D) Effects of the 5’P-gRNA lengths on MhAgo cleavage activity. MhAgo, guides and targets were mixed in a 4:2:1 molar ratio (800 nM MhAgo preloaded with 400 nM guide, plus 200 nM target) and incubated for 30 min at 60 °C, and the reaction buffer contained 5 mM Mn^2+^. The ssDNA and RNA targets are completely complementary to corresponding guides the 5’-end nucleotides of which are C. M indicates synthesized 34-nt ssDNA or RNA product.

### The effect of divalent cations and temperature on cleavage activity of MhAgo

To determine the prerequisites for MhAgo-mediated target cleavage, substrate cleavage was performed in buffer containing different divalent metal ions (Fe^2+^, Co^2+^, Ni^2+^, Cu^2+^, Zn^2+^, Mg^2+^, Ca^2+^ and Mn^2+^). The results showed that MhAgo had the highest endonuclease activity in the presence of Mn^2+^ when using all four types of guides (Supplementary Figure S2A-D). Except for Mn^2+^, MhAgo can utilize Mg^2+^ as cations to cleave DNA when guided by 5’OH and 5’P-RNA (Supplementary Figure S2B and D, upper panel). Besides, MhAgo was able to utilize Co^2+^ and Ni^2+^ as cations to mediate 5’OH-RNA guided DNA target cleavage (Supplementary Figure S2B, upper panel). Meanwhile, MhAgo can also catalyze RNA target cleavage with 5’OH-gDNA or 5’OH-gRNA in the presence of Co^2+^ (Supplementary Figure S2A and B, lower panel). Then we studied the effect of different concentrations of Mg^2+^ or Mn^2+^ on the activity of MhAgo. The results revealed that the higher the Mn^2+^ concentration between 0.2 and 10 mM, the better cleavage efficiency of MhAgo (Supplementary Figure S2E-H). In addition, titration of Mg^2+^ ions showed that MhAgo was active between 1 and 10 mM Mg^2+^ when using gRNA to target ssDNA (Supplementary Figure S2F and H, upper panel). But we did not observe gDNA mediated target cleavage and 5’OH-gRNA mediated RNA cleavage even if the Mg^2+^ concentration was increased to 10 mM (Supplementary Figure S2E-H). When targeting RNA, the lower the concentration of divalent ions, the stronger the non-specific activity of MhAgo (Supplementary Figure S2E-H, lower panel). We guess that the binding of divalent ions may reduce the non-specific activity of MhAgo. The cleavage efficiency of MhAgo was comparable in buffer containing 5 mM Mn^2+^ and 10 mM Mn^2+^. Thus, we performed cleavage assays in buffer containing 5 mM Mn^2+^ in most experiments.

To further explore the effect of temperature on cleavage activity of MhAgo, we performed cleavage assays at different temperatures ranging from 30 to 75 °C. The activity of ssDNA cleavage mediated by 5’OH-gDNA, 5’P-gDNA and 5’P-gRNA was enhanced with temperature increasing from 45 to 65 °C, and rapidly decreased at 70 °C (Figure 3A, Supplementary Figure S3A, C and D, upper panel). When using 5’OH-gRNA to target ssDNA, MhAgo was active in a much wider range of temperature between 37 to 70 °C and had higher endonuclease activity at 60-70 °C (Figure 3A, Supplementary Figure S3B, upper panel). For RNA targets cleavage, the activity of MhAgo was enhanced with temperature increasing from 37 to 60 °C directed by all four types of guides (Figure 3A, Supplementary Figure S3A-D, lower panel). Distinct to a highly stable DNA target, the RNA target may be degraded partially at 65 °C or higher temperature. Thus, we performed cleavage assays at 60 °C in most experiments.

**Figure 3.**
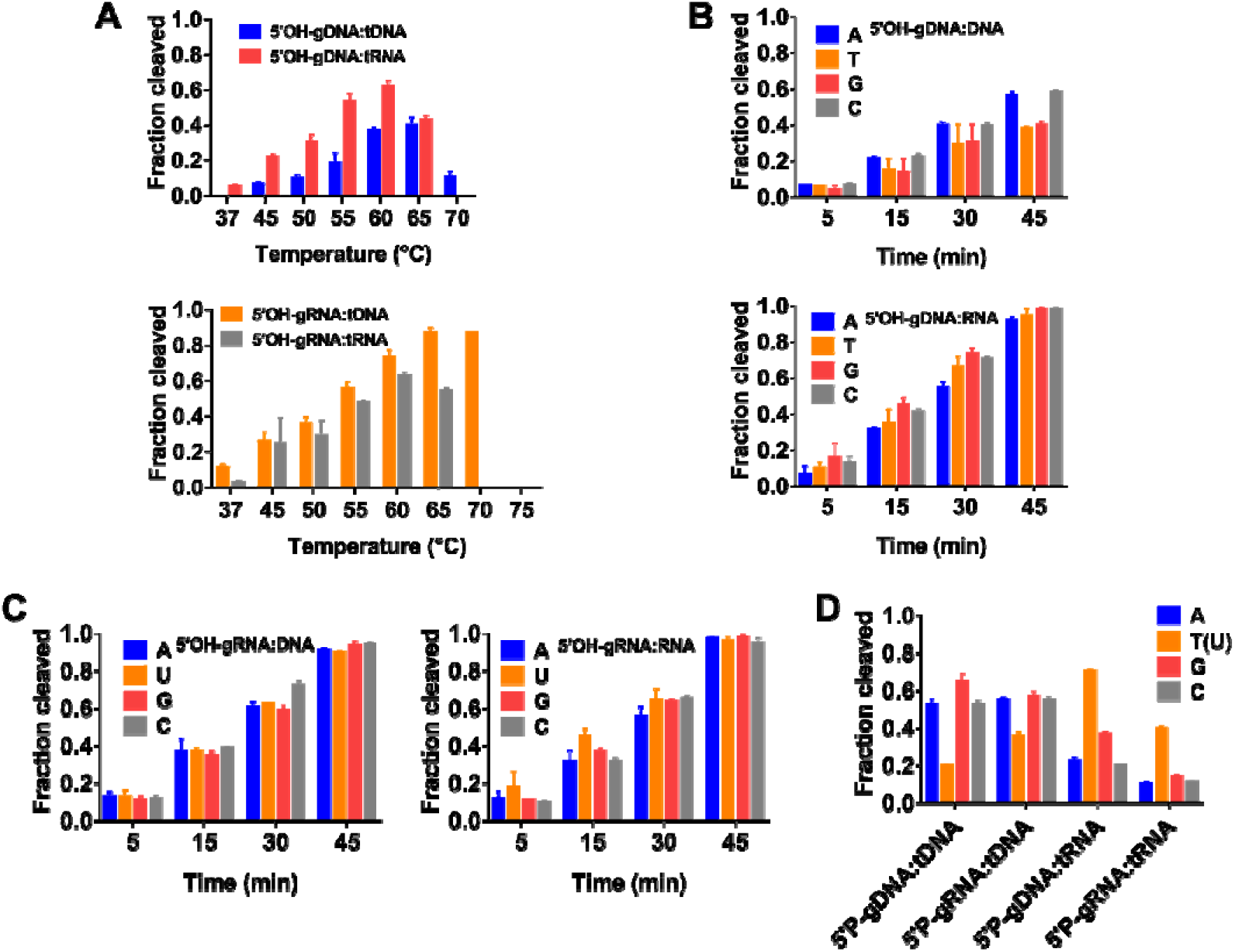
Effects of temperatures and 5’-end nucleotides of guides on MhAgo activity. (A) Effects of temperatures on MhAgo activity mediated by 5’OH-guides. (B) Effects of the 5’-end nucleotides of 5’OH-gDNA on cleavage activity. (C) Effects of the 5’-end nucleotides of 5’OH-gRNA on cleavage activity. (D) Effects of the 5’-end nucleotides of 5’P-guides on cleavage activity. MhAgo, guides and targets were mixed in a 4:2:1 molar ratio (800 nM MhAgo preloaded with 400 nM guide, plus 200 nM target) and the reaction buffer contained 5 mM Mn^2+^. The reaction time in (A) and (D) was 30 min. The reaction temperature in (B), (C) and (D) was 60 °C. Data are represented as the mean ± SD from three independent experiments. In (A), the ssDNA targets are completely complementary to corresponding guides the 5’-end nucleotides of which are C and the RNA targets are completely complementary to corresponding guides the 5’-end nucleotides of which are T or U.

### Effects of 5’-end nucleotides on target cleavage

Many Agos are reported to have preferences for the first nucleotide in the guide strand (18,28). To investigate the effect of 5’-end nucleotides of guides on the cleavage activity of MhAgo, we designed DNA and RNA guides variants with different 5’-terminal nucleotides but otherwise identical sequences. We first tested MhAgo mediated target cleavage activity guided by 5’OH-DNA with different 5’-terminal nucleotides. The results showed that MhAgo mediated DNA cleavage activity was better when MhAgo was loaded with 5’OH-gDNA containing 5’-A and 5’-C compared with 5’-T and 5’-G (Figure 3B, Supplementary Figure S4A, upper panel). And, we did not find an obvious 5’-end nucleotides preference when using 5’OH-gDNA to target RNA (Figure 3B, Supplementary Figure S4A, lower panel). Then we tested whether MhAgo had preferences for a specific 5’-end nucleotides of 5’OH-gRNA. According to our experiments, whether targeting ssDNA or RNA, MhAgo had no obvious preference for the 5’-end nucleotides of 5’OH-gRNA (Figure 3C, Supplementary Figure S4B). Next, we tested MhAgo mediated target cleavage activity guided by 5’P-DNA and RNA with different 5’-terminal nucleotides. For 5’P-gDNA and 5’P-gRNA mediated DNA cleavage, MhAgo had better activity when the 5’-end nucleotides were A, G and C (Figure 3D, Supplementary Figure S4C, left panel). For 5’P-gDNA and 5’P-gRNA mediated RNA cleavage, the strongest activity was observed when the 5’-end nucleotides were T or U (Figure 3D, Supplementary Figure S4C, right panel). The ability of MhAgo to cleave ssDNA and RNA mediated by different lengths and sequence guides makes it possible to become an alternative tool for nucleic acids manipulation.

### Effects of guide-target mismatches on target cleavage

Complementarity between guide sequence and target sequence determines the cleavage specificity of Agos (20,42). To characterize the specificity, we tested the effect of guide-target mismatches on the cleavage activity of MhAgo. It is mentioned that the guide can be divided into several regions, including the 5’-end region (position 1), the 5’-seed region (positions 2-8), the central region (positions 9-12), the 3’-supplementary region (positions 13-15) and 3’-tail region (positions 16-18) (20). We first analyzed the effects of single mismatches between 5’OH-gDNA and target strands on MhAgo activity. Owing to the activity of 5’OH-gDNA mediated DNA cleavage was weaker, we performed the cleavage assay at 60 °C for 1 h. We found that single mismatches in the 5’-end region of 5’OH-gDNA had no effect on the cleavage efficiency of MhAgo, and single mismatches in the 3’-tail region of 5’OH-gDNA reduced the target cleavage activity of MhAgo, whereas single mismatches that occurring in the 5’-seed (except for position 2), 3’-supplementary and central regions of 5’OH-gDNA showed a drastic reduction in ssDNA and RNA cleavage efficiency (Figure 4A and B, Supplementary Figure S5A). Despite single mismatches in the seed region, 5’-end and tail regions of 5’OH-gRNA had little effect on MhAgo mediated target cleavage, single mismatches in the central and 3’-supplementary regions resulted in a strong decrease in cleavage efficiency (Figure 4C and D, Supplementary Figure S5B). Furthermore, we analyzed the effects of single mismatches between 5’P-guides and target strands on MhAgo activity. When guided by 5’P-DNA, single mismatches in the 5’-end and 3’-tail regions of guides had little effect on MhAgo activity and single mismatches in the 5’-seed (except for position 2), central, 3’-supplementary regions of guides showed a drastic reduction in cleavage efficiency (Supplementary Figure S5C). When guided by 5’P-RNA, single mismatches in the 5’-end, 5’-seed and 3’-tail regions of guides had no substantial decrease on MhAgo activity and single mismatches in the central and 3’-supplementary regions of guides showed a obvious reduction in target cleavage efficiency (Supplementary Figure S5D). In summary, single mismatches occurring in different regions of guides had various effects on cleavage efficiency, depending on the types of guides. Importantly, whether it was gDNA or gRNA, single mismatches that occurring in the central and 3’-supplementary regions of guides would greatly reduce the cleavage efficiency. The single mismatches in the 12th nucleotide counting from the 5’ end of guides completely abolished MhAgo activity. The ability to distinguish single bases makes MhAgo possible for nucleic acid detection.

**Figure 4.**
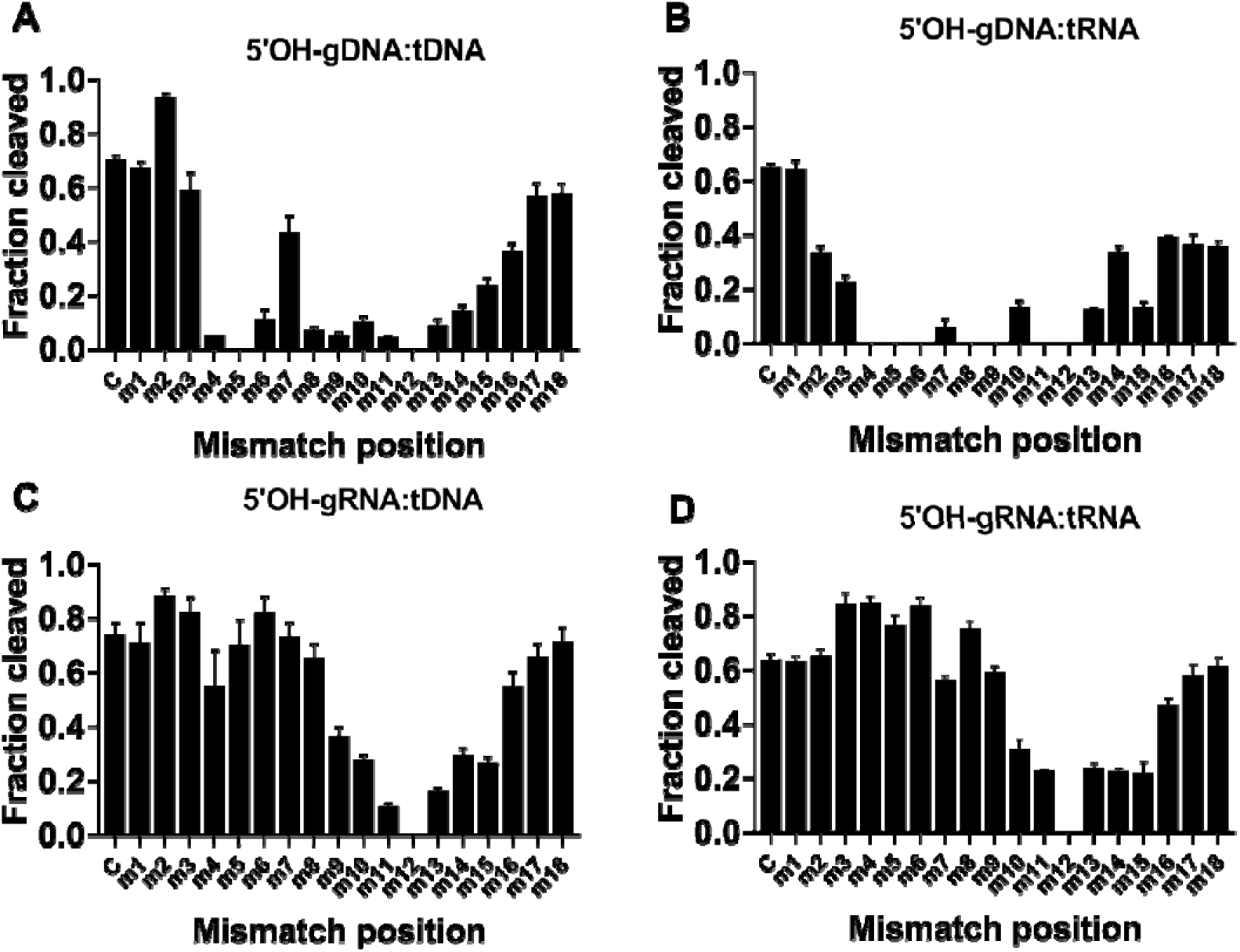
Effects of guide-target mismatches on MhAgo activity. (A) Effects of guide-target mismatches on DNA cleavage activity mediated by 5’OH-gDNA. (B) Effects of guide-target mismatches on RNA cleavage activity mediated by 5’OH-gDNA. (C) Effects of guide-target mismatches on DNA cleavage activity mediated by 5’OH-gRNA. (D) Effects of guide-target mismatches on RNA cleavage activity mediated by 5’OH-gRNA. All experiments were performed at the 4:2:1 MhAgo:guide:target molar ratio in reaction buffer containing 5 mM Mn^2+^ ions at 60 °C. The assay in (A) was performed for 1 h. The assay in (B), (C) and (D) was performed for 30 min. Data are represented as the mean ± SD from three independent experiments. The ssDNA and RNA targets are completely complementary to corresponding guides the 5’-end nucleotides of which are C.

### The efficiency of different guides and targets binding and cleavage

To further explore the cleavage activity of MhAgo with these optimized cleavage conditions, we measured the efficiency of target cleavage using either 5’OH or 5’P-guides. For DNA targets, all four types of guides were able to direct efficiently cleavage, with only slightly lower efficiency with 5’OH-gDNA (Figure 5A, Supplementary Figure S6). For RNA targets, the cleavage efficiency of MhAgo with 5’OH-gDNA, 5’P-gDNA and 5’OH-gRNA was comparable. The weakest RNA cleavage efficiency was observed with 5’P-gRNA (Figure 5B, Supplementary Figure S6).

**Figure 5.**
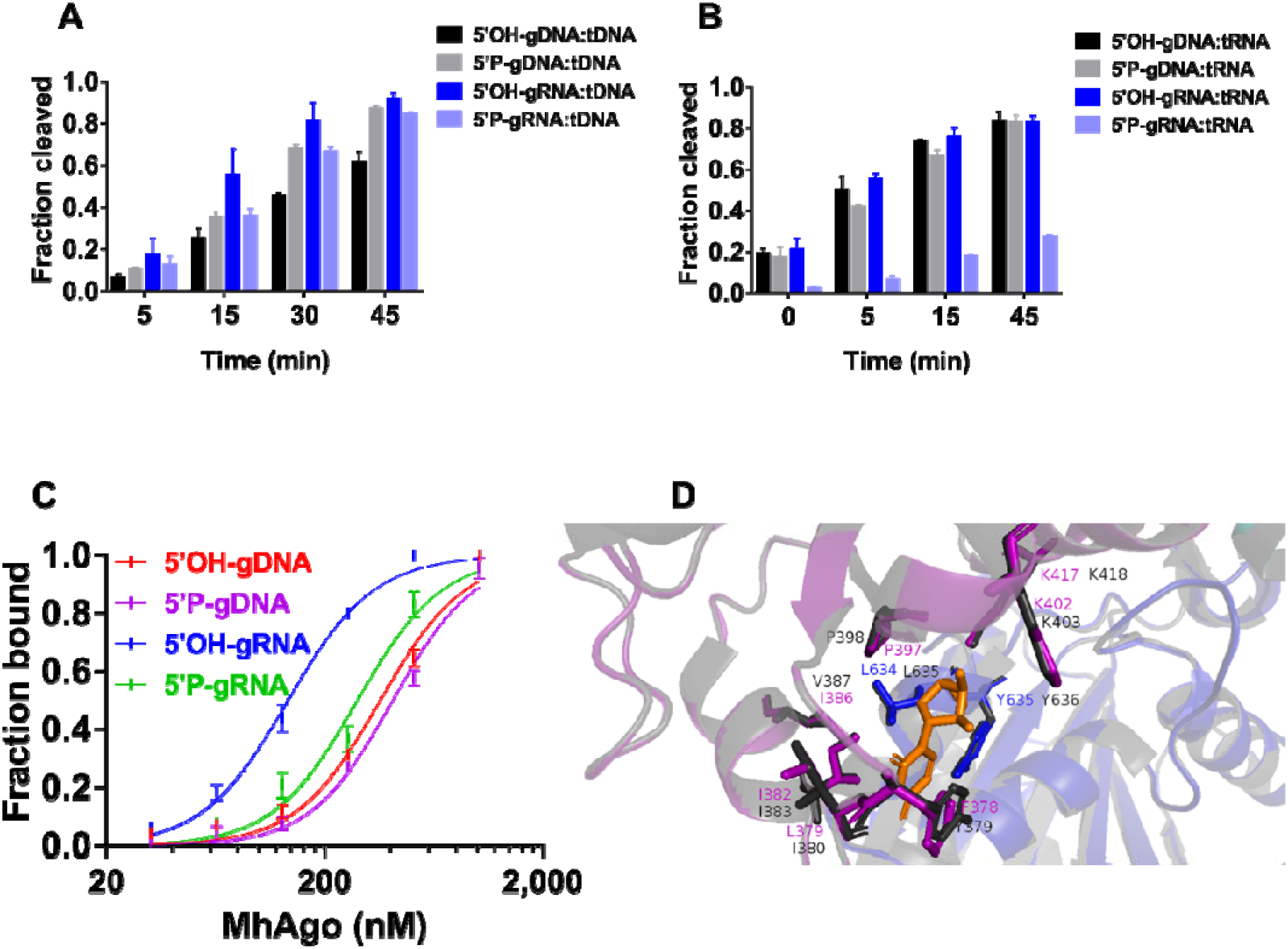
The analysis of different guides and targets binding and cleavage. (A) Cleavage analysis of ssDNA targets using different guides in a time course experiment. (B) Cleavage analysis of RNA targets using different guides in a time course experiment. The cleavage experiments were performed at the 4:2:1 MhAgo:guide:target molar ratio in reaction buffer containing 5 mM Mn^2+^ ions at 60 °C. (C) Binding analysis of different guides including 5’OH-gDNA, 5’P-gDNA, 5’OH-gRNA and 5’P-gRNA by MhAgo. The fraction of bound guide is plotted against protein concentration and fitted using the model of specific binding with the Hill slope. MhAgo binds the 5’OH-gDNA and 5’P-gDNA with average K_d_ values of 359.50 ± 31.39 nM and 410.30 ± 31.74 nM, respectively. MhAgo binds the 5’OH-gRNA and 5’P-gRNA with average K_d_ values of 136.00 ± 4.90 nM and 282.70 ± 24.93 nM, respectively. Data are represented as the mean ± SD from three independent experiments. (D) The MID (purple) and PIWI (blue) domains of simulated MhAgo and MpAgo (grey) structures form the binding pocket for the 5’ end of the RNA guide. The amino acid residues in the MID domain of MhAgo are colored purple; the amino acid residues in the PIWI domain of MhAgo are colored blue; the amino acid residues of MpAgo are colored black. In (A) and (B), the ssDNA targets are completely complementary to corresponding guides the 5’-end nucleotides of which are C and the RNA targets are completely complementary to corresponding guides the 5’-end nucleotides of which are T or U. The 5’-end nucleotides of guides used in EMSA are C.

To explore the affinity of MhAgo to different types of guides, we performed Electrophoretic mobility shift assay (EMSA) and measured the dissociation constants (K_d_) for different guides binding by MhAgo. As shown, MhAgo had similar affinity for 5’P-gDNA (K_d_: 410.30 ± 31.74 nM) and 5’OH-gDNA (K_d_: 359.50 ± 31.39 nM) (Figure 5C, Supplementary Figure S7A and B). This can be the reason why MhAgo can utilize both 5’OH-gDNA and 5’P-gDNA to cleave targets. MhAgo had the strongest affinity for 5’OH-gRNA (K_d_: 136.00 ± 4.90 nM), followed by 5’P-gRNA (Kd: 282.70 ± 24.93 nM) (Figure 5C, Supplementary Figure S7C and D). This roughly explained why MhAgo performed the best cleavage activity guided by 5’OH-RNA.

MhAgo shared a high sequence identity (65.41%) with MpAgo. To shed light on the molecular mechanism of the MhAgo function, we obtained the structure of MhAgo according to homology modeling with MpAgo/gRNA complex as a template (Supplementary Figure S8A). The MID binding pocket of MpAgo is lined with hydrophobic residues (I383, V387, P398, L635, Y636), which surround the 5’ hydroxyl group of the guide strand (27). According to the structure-based sequence alignment of MhAgo and MpAgo, the corresponding amino acid residues in MhAgo are I382, I386, P397, L634 and Y635. The first nucleotide of the guide has stacking interactions with Y379 and I380 in the MID binding pocket of MpAgo (27) and the corresponding amino acid residues in MhAgo are F378 and L379 (Figure 5D, Supplementary Figure S8B). The residues V387, Y379 and I380 of MpAgo are different from the residues I386, F378 and L379 of MhAgo. To explore whether the three amino acid residues resulted in different properties of MhAgo and MpAgo, we mutated the three residues separately to obtain mutant_I386V/A, mutant_F378Y/A, mutant_L379I/A and mutant_I386V/F378Y/L379I. The results showed that the seven mutants had little effect on cleavage activity compared to wild-type MhAgo (Supplementary Figure S9). Further work is needed to decipher the underlying molecular mechanisms. The presence of an ordered α-helix (α5) at the C terminus of the PIWI domain is also one of the reasons why MpAgo hinders the binding of 5’P-guides (27). By comparing the structures of TtAgo, MpAgo and MhAgo, we found that MhAgo also had an ordered α5 different from TtAgo at the C terminus of the PIWI domain (Supplementary Figure S8C). The phenomenon may explain why MhAgo preferentially binds to 5’OH-gRNA like MpAgo.

For ssDNA target, all four types of guides were able to direct efficiently cleavage, with only slightly lower efficiency with 5’OH-gDNA (Figure 5A). MhAgo can use 5’OH-gDNA, 5’P-gDNA and 5’OH-gRNA to efficiently cleave RNA and had the weakest RNA cleavage efficiency directed by 5’P-gRNA (Figure 5B). To explain these phenomena, we measured the binding for guide:target duplexes by MhAgo using EMSA. MhAgo had a slightly stronger affinity for 5’OH-guide:target duplexes than 5’P-guide:target duplexes (Figure 6A-D), which may explain why MhAgo had slightly better activity using 5’OH-guides. The affinity of MhAgo for 5’OH-gRNA:tRNA duplex (K_d_: 125.40 ± 6.83 nM) was obviously stronger than for 5’P-gRNA:tRNA duplex (K_d_: 231.9 ± 5.07 nM) (Figure 6D), which may be one of the reasons that MhAgo was only able to cleave RNA target with low efficiency guided by 5’P-RNA. MhAgo performed stronger affinity for 5’P-gRNA:tDNA duplex (K_d_: 147.50 ± 4.52 nM) than for 5’P-gRNA:tRNA (K_d_: 231.9 ± 5.07 nM) (Figure 6E), which may explain why MhAgo can utilize 5’P-gRNA more efficiently cleave DNA than cleave RNA. Formation of the MhAgo–duplex complexes and the disappearance of free strands were observed in all eight combinations of guides and targets (Supplementary Figure S10). These observations further illustrated the eight cleavage patterns of MhAgo and we suggested that the binding activity of MhAgo may influence its cleavage activity.

**Figure 6.**
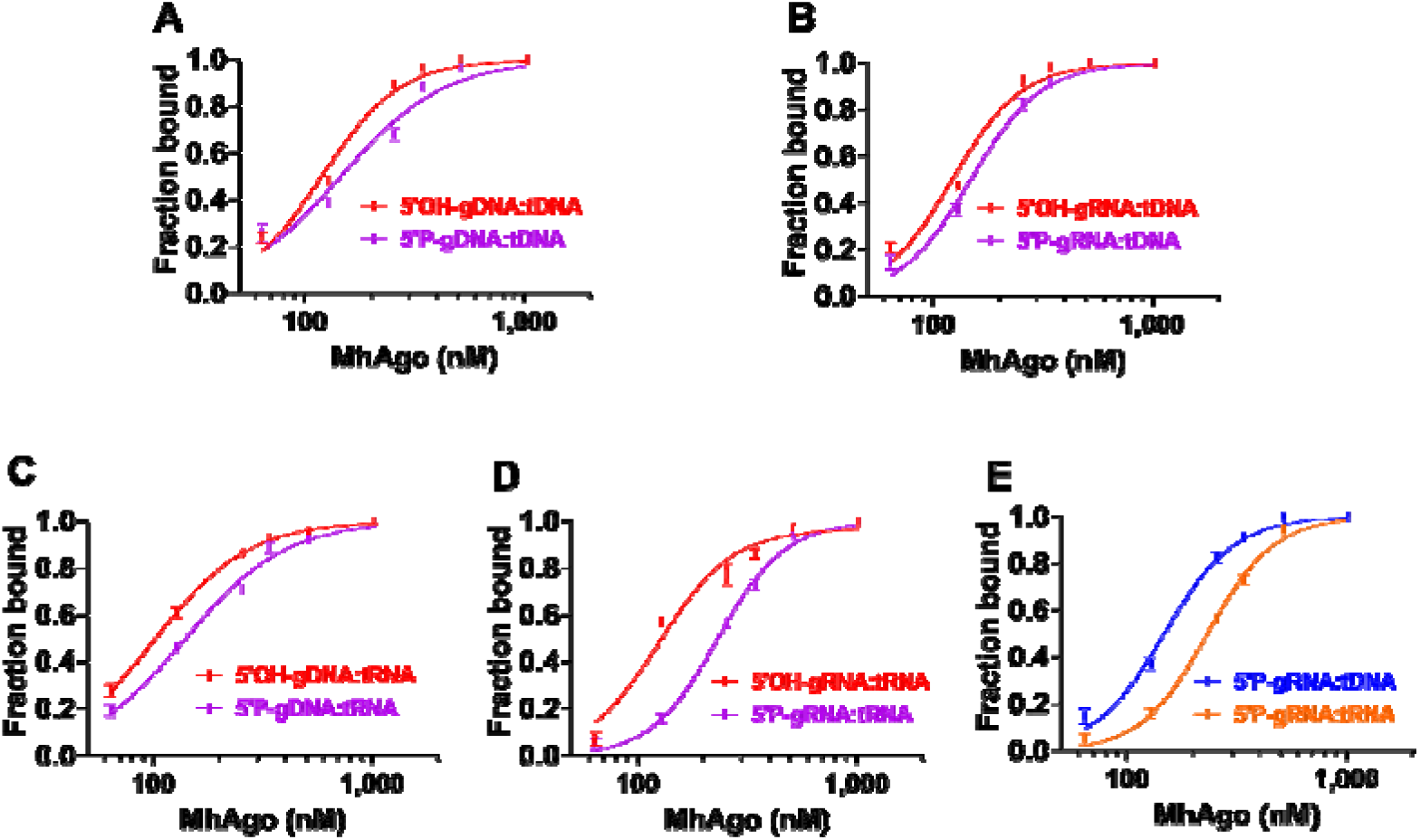
Binding analysis of different guide:target duplexes by MhAgo. (A) Binding of gDNA:tDNA duplexes by MhAgo. MhAgo binds the 5’OH-gDNA:tDNA duplex and 5’P-gDNA:tDNA duplex with average K_d_ values of 118.90 ± 4.80 nM and 144.80 ± 11.51 nM, respectively. (B) Binding of gRNA:tDNA duplexes by MhAgo. MhAgo binds the 5’OH-gRNA:tDNA duplex and 5’P-gRNA:tDNA duplex with average K_d_ values of 121.40 ± 4.66 nM and 147.50 ± 4.52 nM, respectively. (C) Binding of gDNA:tRNA duplexes by MhAgo. MhAgo binds the 5’OH-gDNA:tRNA duplex and 5’P-gDNA:tRNA duplex with average K_d_ values of 102.20 ± 1.69 nM and 140.00 ± 5.37 nM, respectively. (D) Binding of gRNA:tRNA duplexes by MhAgo. MhAgo binds the 5’OH-gRNA:tRNA duplex and 5’P-gRNA:tRNA duplex with average K_d_ values of 125.40 ± 6.83 nM and 231.9 ± 5.07 nM, respectively. (E) Binding of 5’P-gRNA:target duplexes by MhAgo. The fraction of bound guide is plotted against protein concentration and fitted using the model of specific binding with the Hill slope. MhAgo_DM protein was added after 64 nM guides and 64 nM targets were annealed to form duplexes. Then reaction mixtures were incubated for 30 min at 55 °C. Non-labeled ssDNA and RNA targets are used in EMSA. The ssDNA and RNA targets are completely complementary to corresponding guides the 5’-end nucleotides of which are C.

## DISCUSSION

RNA interference pathways in eukaryotes use small single-stranded RNA molecules that guide eAgos to target complementary RNA targets (4). The *in vivo* functions of pAgos are still unclear. Some studies demonstrate that pAgos function in host defense by DNA-guided DNA interference (5,8,15). However, the discovery of RsAgo and MpAgo brings new insights into the understanding of pAgos. RsAgo specifically recognizes 5’P-RNA as the guide strand and DNA as the target strand (9,28). MpAgo prefers 5’OH-gRNA that are chemically distinct from the 5’P-gDNA used by other pAgos studied to date (27,39). Here, we reported a novel Argonaute protein from *Marinitoga hydrogenitolerans* (MhAgo) which was different from previously reported pAgos. MhAgo can utilize all types of guides (5’OH-gDNA, 5’P-gDNA, 5’OH-gRNA and 5’P-gRNA) to cleave complementary ssDNA and RNA targets.

Most pAgos use divalent cation for interaction with the phosphate group of 5’-end of 5’P-guides. In contrast to other pAgos, MpAgo binds unphosphorylated guides and has a more hydrophobic binding pocket without metal ions. But it is vital for the catalytic activity of pAgos that DEDX in PIWI domain chelate catalytic divalent metal ions, Mg^2+^ or Mn^2+^ (43). Like most Agos, MhAgo also needed divalent metal ions for cleavage. In the presence of Mn^2+^, MhAgo exhibited all eight cleavage activities. While, in presence of Mg^2+^, MhAgo only had gRNA-mediated DNA cleavage activity. MhAgo had high activity at temperatures ranging from 45 °C to 65 °C. Meanwhile, MhAgo also had weak activity at 37 °C. In the future, we may also mine such versatile mesophilic Agos with potentials for DNA/RNA editing. MhAgo can cleave target ssDNA and RNA mediated by guides longer than 12 nt and efficiently cleave target with16-21 long guides, which was similar to most pAgos (16,20). The limitation of guide length makes it possible for nucleic acid detection, such as PfAgo (35). For some Agos, clear preferences and a high abundance of guide strands with particular 5’-nucleotides are found. When MjAgo uses gDNA to cleave ssDNA target, the cleavage efficiency is significantly higher using a guide with a 5’-purine than with a 5’-pyrimidine (18). When KpAgo uses gDNA to cleave RNA target, guides beginning with 5’-T, A or C cleave the target comparably (~60%) whereas guides with a 5’G display only ~40% cleavage (44). When RsAgo uses gRNA to recognize DNA target, the MID domain has the highest affinity for 5’U-gRNA (28). When hAgo2 uses gRNA to recognize RNA target, the MID domain has the highest affinity for UMP followed by AMP, whereas the affinities for CMP and GMP are considerably weaker (45). When using 5’OH-gDNA to target complementary ssDNA, MhAgo had a slightly higher efficiency of target cleavage with A and C as the 5’ terminal nucleotides. And, when using 5’OH-gDNA to target RNA and using 5’OH-gRNA to target ssDNA/RNA, MhAgo had no obvious preference for the 5’-end nucleotides of guides. For DNA targets, MhAgo had the weakest activity using T or U as the 5’ end of 5’P-guides. For RNA targets, MhAgo had the strongest activity using T or U as the 5’ end of 5’P-guides.

Previous studies of eAgos and several pAgos show that mismatches between the guide and target strands may have large effects on the target recognition. When LrAgo uses gDNA to cleave ssDNA target, single mismatches surrounding the cleavage site and in the 3’-supplementary region of guides strongly decrease the efficiency of target cleavage (20). When TtAgo uses gDNA to cleave RNA target, cleavage is either abolished (positions 9 and 10) or reduced (positions 11 and 12) by mismatches surrounding the cleavage site and single mismatches in the seed region (position 2–8) show reduced cleavage activity (46). When guided by RNA, mismatches in the seed region have a large effect on the efficiency of target recognition or cleavage, such as RsAgo, MpAgo, AfAgo, Mouse Ago2 (40,47–49). When guided by DNA, MhAgo can tolerate guide/target mismatches in the 5’-end and 3’-tail regions of guides but were sensitive to mismatches in the 5’-seed, central and 3’-supplementary regions of guides. When guided by RNA, MhAgo can tolerate guide/target mismatches in the 5’-end, 5’-seed and 3’-tail regions of guides but were sensitive to mismatches in the central and 3’-supplementary regions of guides. These results showed that MhAgo had excellent specificity in all the eight cleavage patterns. By comparing the cleavage efficiency of ssDNA and RNA targets directed by different guides, we found that MhAgo can more efficiently cleave ssDNA guided by 5’P-gDNA, 5’OH-gRNA and 5’P-gRNA, and more efficiently cleave RNA guided by 5’OH-gDNA, 5’P-gDNA, and 5’OH-gRNA. The omnipotence and specificity of MhAgo make it have a wider application prospect.

The distinctive characteristics of MhAgo arise our interest to explore its molecular mechanisms. We performed EMSA to explore the affinity of MhAgo to different guides and found that MhAgo had the highest affinity for 5’OH-gRNA. This was consistent with that MhAgo performed the best cleavage activity guided by 5’OH-RNA. We also performed EMSA to explore the affinity of MhAgo to different guide:target duplexes and found that MhAgo was able to bind all eight nucleic acid duplexes, which roughly explained why MhAgo exhibited all eight patterns of cleavage. MhAgo had the weakest affinity for 5’P-gRNA:RNA target duplex. This was consistent with that MhAgo performed the weakest RNA cleavage activity guided by 5’P-RNA, which suggested that the cleavage activity of MhAgo was related to its binding activity. MhAgo shared high-level sequence identity with MpAgo. Thus, we used MpAgo as a template to perform homology modeling analysis of MhAgo. We found that the predicted MID binding pocket of MhAgo was a little wider than that of MpAgo (Supplementary Figure S8D), which may explain the ability of MhAgo to utilize all four types of guides for cleavage. We mutated several amino acids which may involve in the formation of MID pockets of MhAgo into several amino acids corresponding MpAgo (27). These mutants had little effect on cleavage activity compared to wild-type MhAgo. This may be because the guides and targets binding by MhAgo involves the function of other amino acids residues and protein conformation different from other Agos. These require further biochemical experiments and structural analysis to explain.

Like eAgos, most pAgos interact with 5’P-guides, but in contrast to eAgos, most pAgos have a higher affinity for DNA guides than RNA guides (16,20). Some eAgos also use 5’OH-gRNA to target complementary RNA *in vitro* (42). The pAgos are much more diverse than eAgos in the case of *in vitro* function though they have high structural homology. AfAgo shares significant sequence homology with human Argonaute proteins and can utilize RNA guides to recognize RNA targets (12,29). AaAgo has RNA cleavage activity and has better activity guided by 5’P-gDNA than by 5’OH-gDNA (26). MpAgo has preferences for 5’OH-gRNA and cleaves DNA targets better than RNA targets (27). RsAgo only uses 5’P-gRNA to recognize DNA targets and has no catalytic activity (28). CpAgo can cleave both DNA and RNA targets guided by 5’OH and 5’P-gDNA (21). The recently reported KmAgo is the most versatile pAgo known so far (23,24) (Supplementary, Table S1). Here, the MhAgo possessed all the above special activities. Phylogenetic analyses indicate that the evolutionary journey of the Agos starts in prokaryotes and long pAgos from several euryarchaeal species, mainly thermophiles, are grouped with eAgos (1,2). We guess that the most primitive pAgo has all the cleavage activity and is thermophilic. As the species evolves, part of the activity is lost or weakened, which constitutes the current various Agos. The discovery of MhAgo expands our knowledge of Agos, and we need further exploration to understand the evolutionary journey of Agos.

## SUPPLEMENTARY DATA

Supplementary Data are available at NAR Online.

