## Supplemental Figure 1-10, Table 1-2 for "A bacterial Argonaute with efficient DNA and RNA cleavage activity guided by small DNA and RNA"

**TABLE AND FIGURES LEGENDS**

**A**

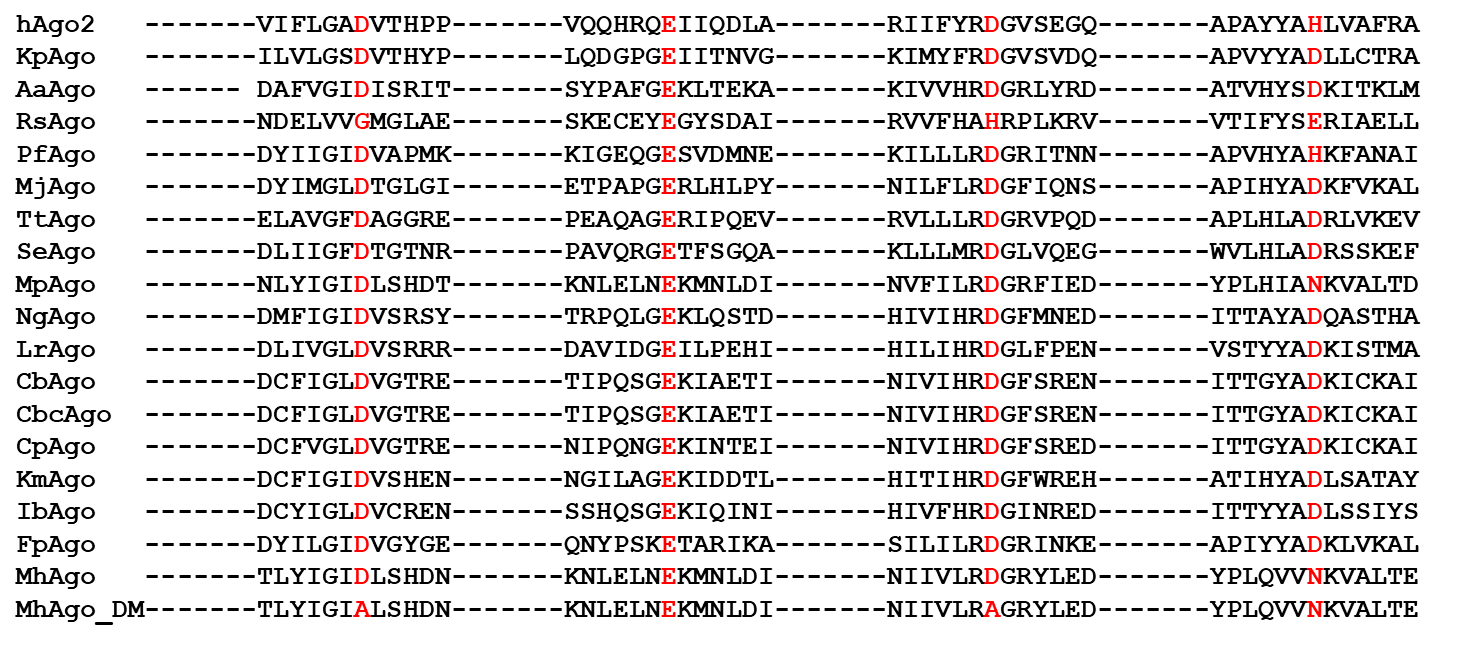

**B**

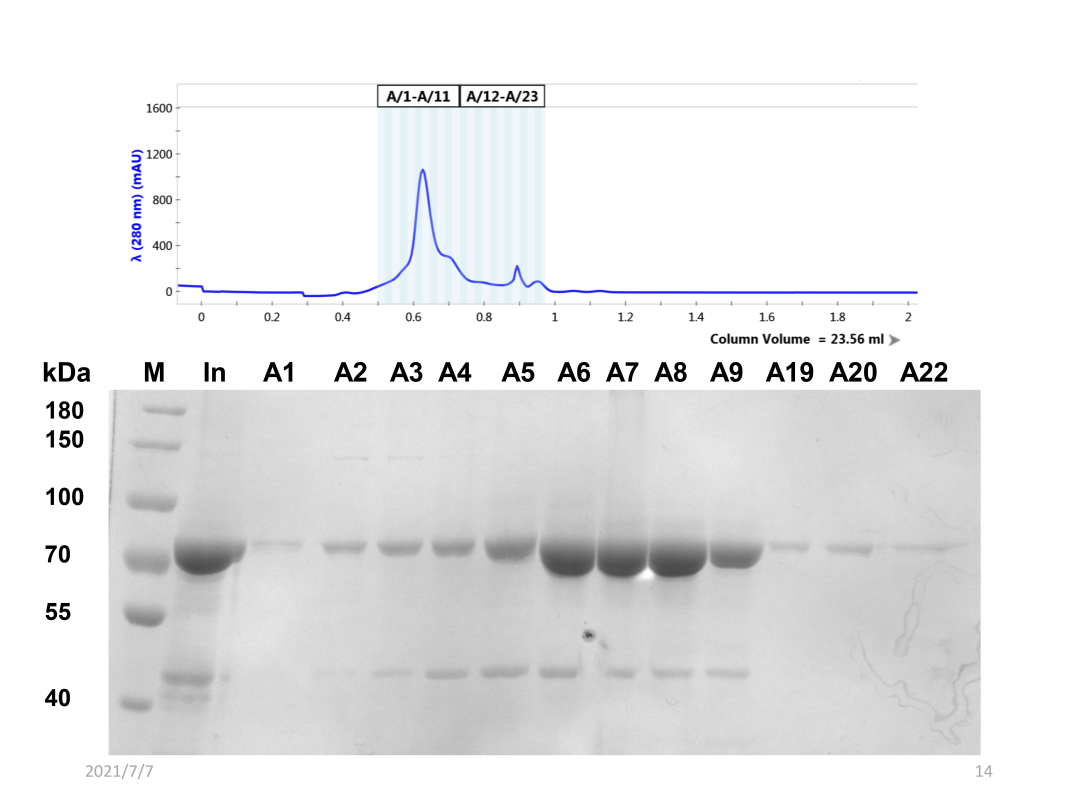

**C**	**D**

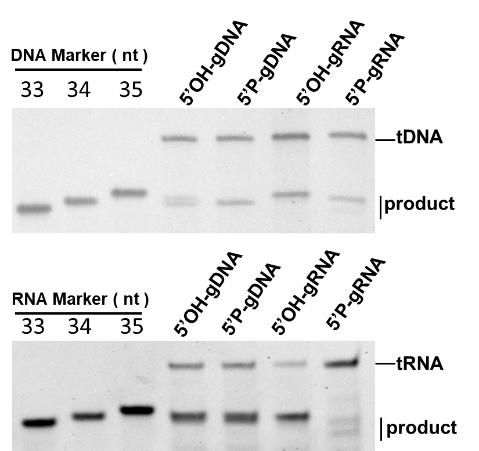

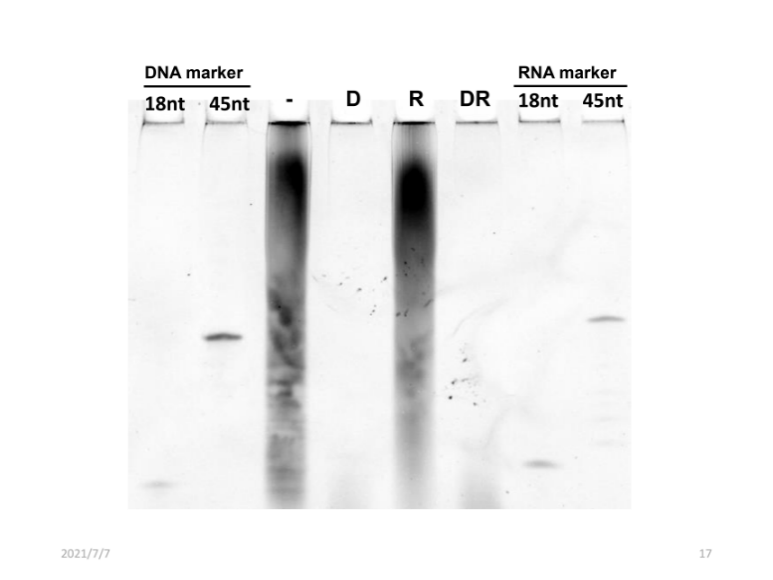

Supplementary Figure S1. DEDN catalytic site of MhAgo. (A) Multiple sequence alignment of conserved amino acid residues (marked by red) of the DEDX tetrad localized in the PIWI domain of Agos. The catalytically dead variant of MhAgo (MhAgo_DM) with two amino acid substitutions within the catalytic tetrad is also shown. (B) Size-exclusion chromatogram and SDS/PAGE of wild-type MhAgo. Proteins are purified using a Superdex 200 16/60 (GE Life Sciences) size exclusion column. (C) Determining the DNA (upper panel) and RNA (lower panel) cleavage site with different guides. DNA marker (33, 34, 35 nt) and RNA marker (33, 34, 35 nt) are chemically synthesized. (D) Nucleic acids that co-purified with MhAgo are treated with either RNAse A, DNAse I or both, and are analyzed by denaturing polyacrylamide gel electrophoresis. In (C), the ssDNA targets are completely complementary to corresponding guides the 5’-end nucleotides of which are C and the RNA targets are completely complementary to corresponding guides the 5’-end nucleotides of which are T or U.

**A**	**B**

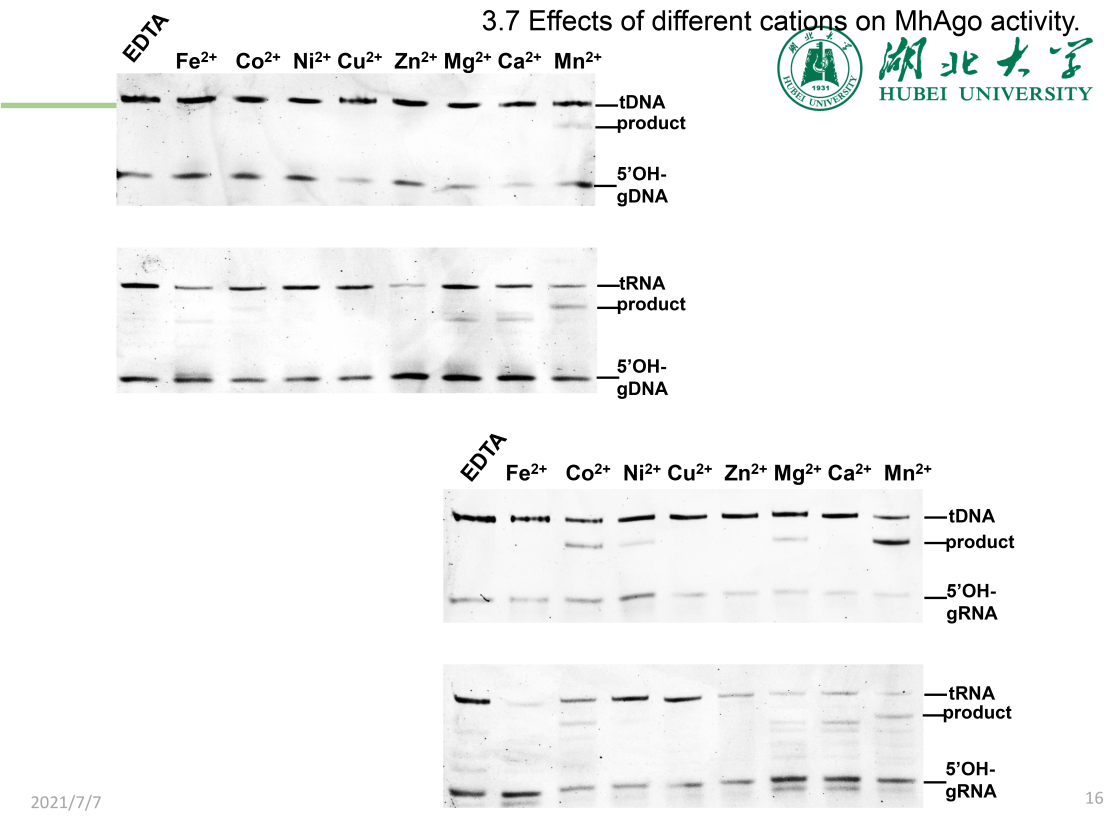

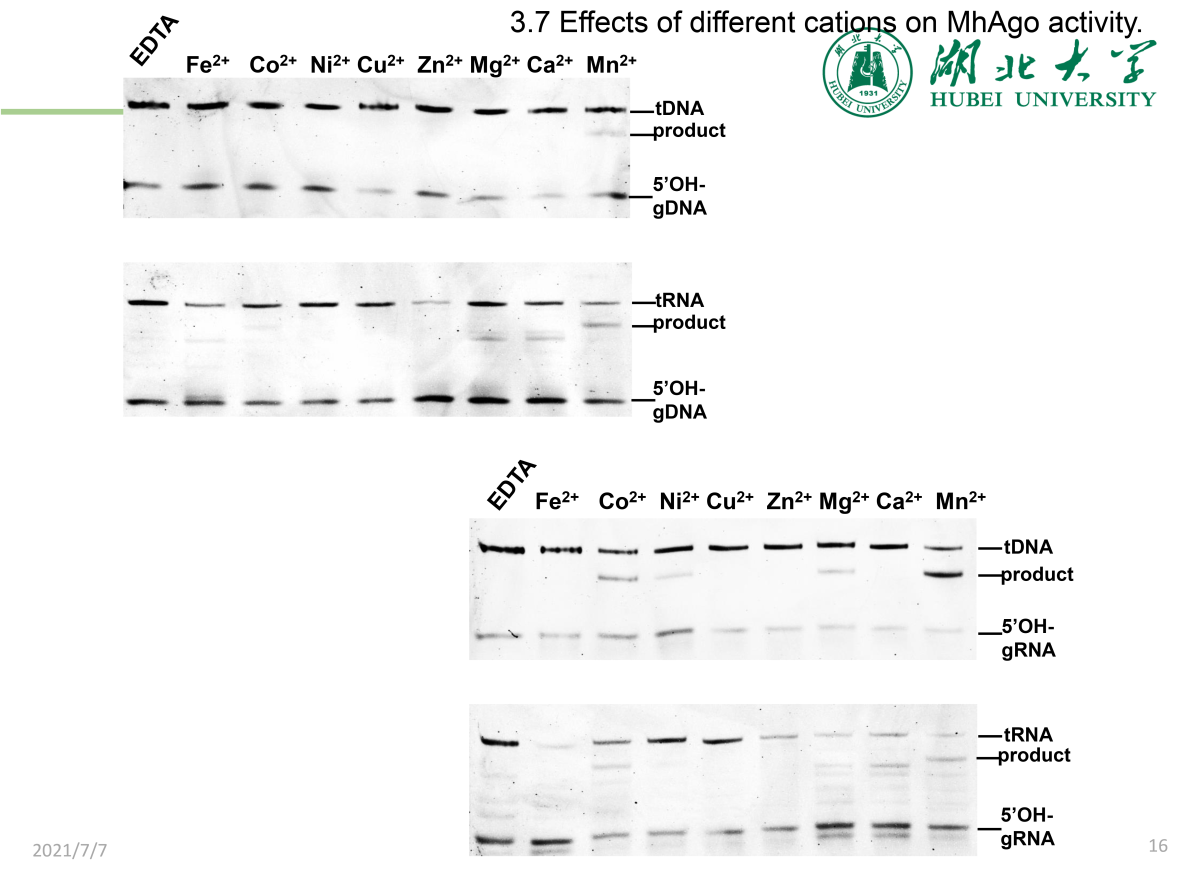

**C	D**

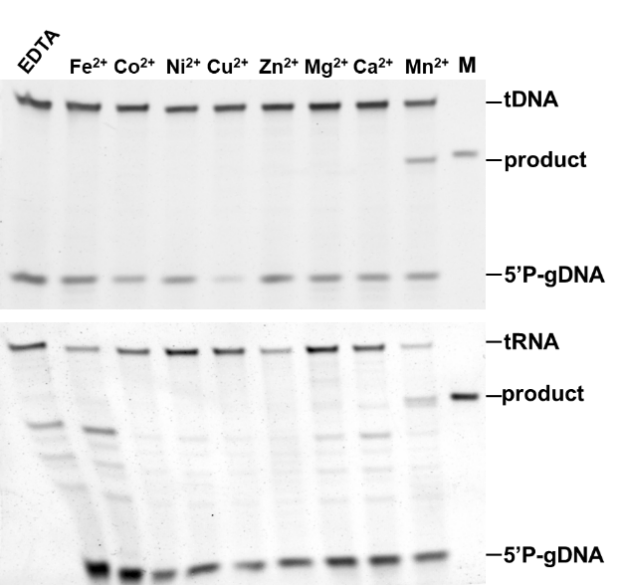

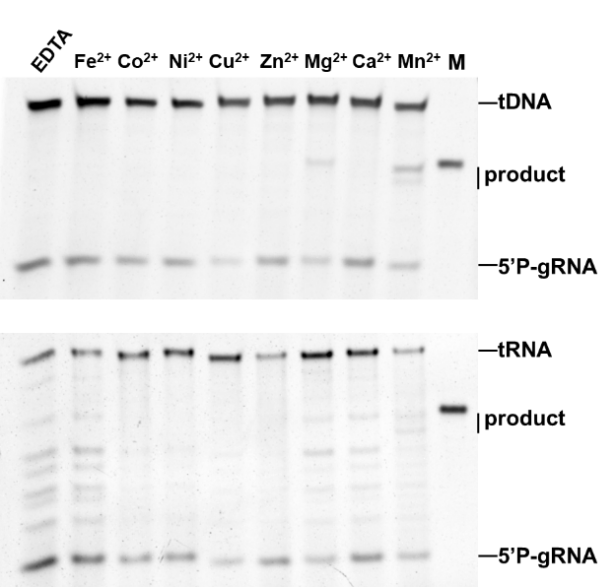

**E**	**F**

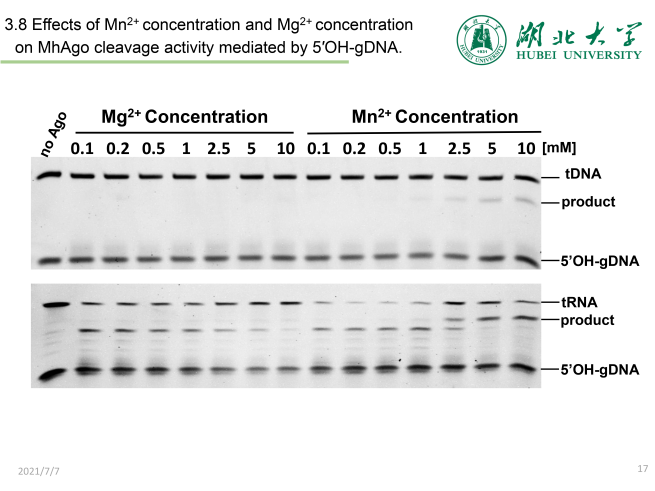

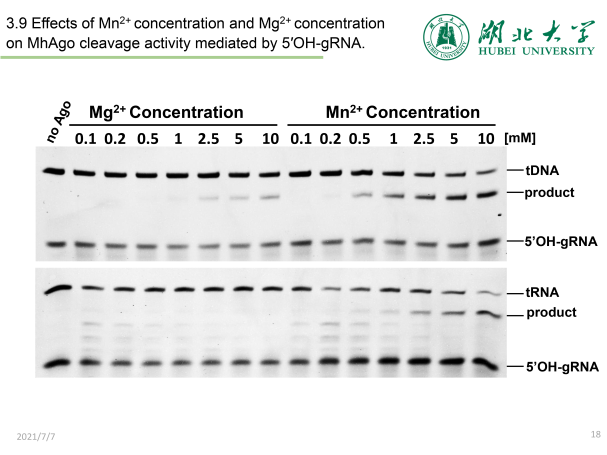

**G	H**

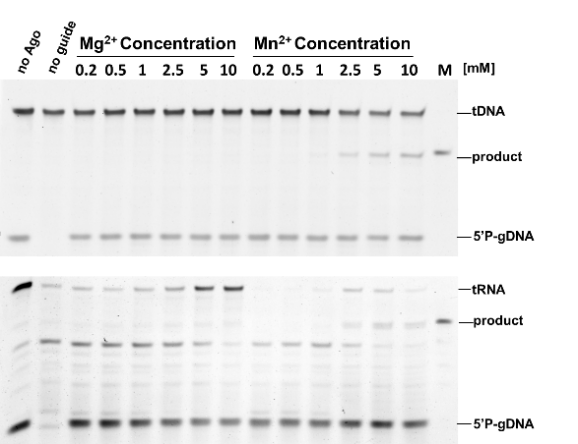

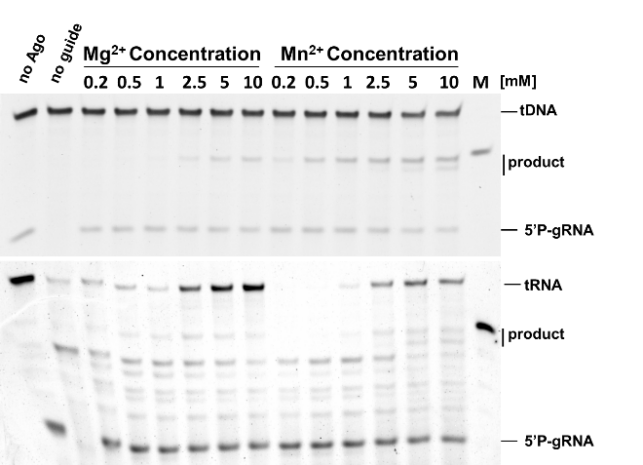

Supplementary Figure S2. Effects of different divalent metal ions on MhAgo activity. (A) Effects of different divalent metal ions on MhAgo activity mediated by 5’OH-gDNA. (B) Effects of different divalent metal ions on MhAgo activity mediated by 5’OH-gRNA. (C) Effects of different divalent metal ions on MhAgo activity mediated by 5’P-gDNA. (D) Effects of different divalent metal ions on MhAgo activity mediated by 5’P-gRNA. (E) Effects of Mn^2+^ concentrations and Mg^2+^ concentrations on target cleavage activity mediated by 5’OH-gDNA. (F) Effects of Mn^2+^ concentrations and Mg^2+^ concentrations on target cleavage activity mediated by 5’OH-gRNA. (G) Effects of Mn^2+^ concentrations and Mg^2+^ concentrations on target cleavage activity mediated by 5’P-gDNA. (H) Effects of Mn^2+^ concentrations and Mg^2+^ concentrations on target cleavage activity mediated by 5’P-gRNA. All reactions were carried out for 30 min at 55 °C with 4:2:1 MhAgo:guide:target molar ratio (800 nM MhAgo preloaded with 400 nM guide, plus 200 nM target). The experiments in (A) and (B) were performed in a reaction buffer containing 5 mM divalent metal ion or EDTA. The ssDNA targets are completely complementary to corresponding guides the 5’-end nucleotides of which are C. The RNA targets are completely complementary to corresponding guides the 5’-end nucleotides of which are T or U. M indicates synthesized 34-nt ssDNA or RNA product.

**A**	**B**

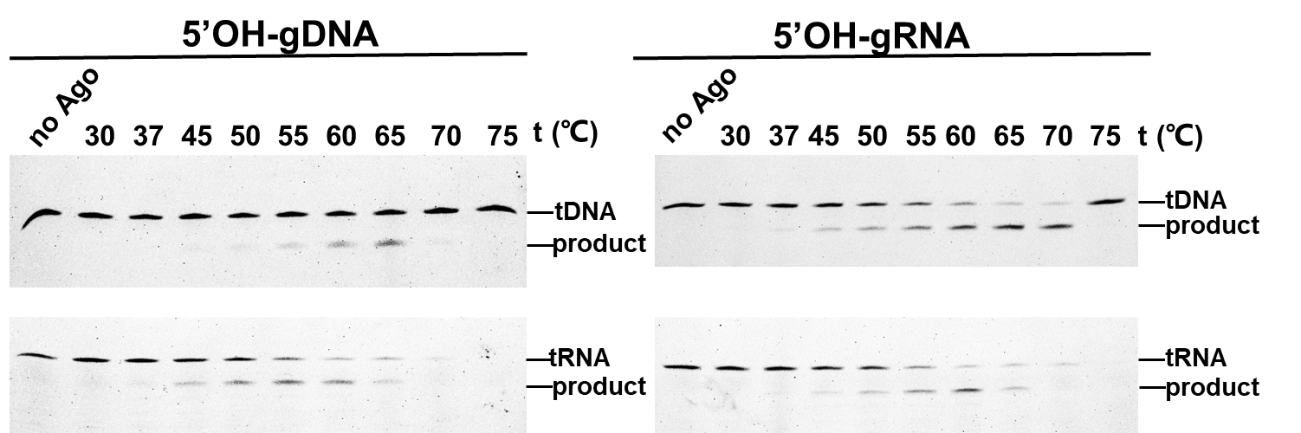

**C	D**

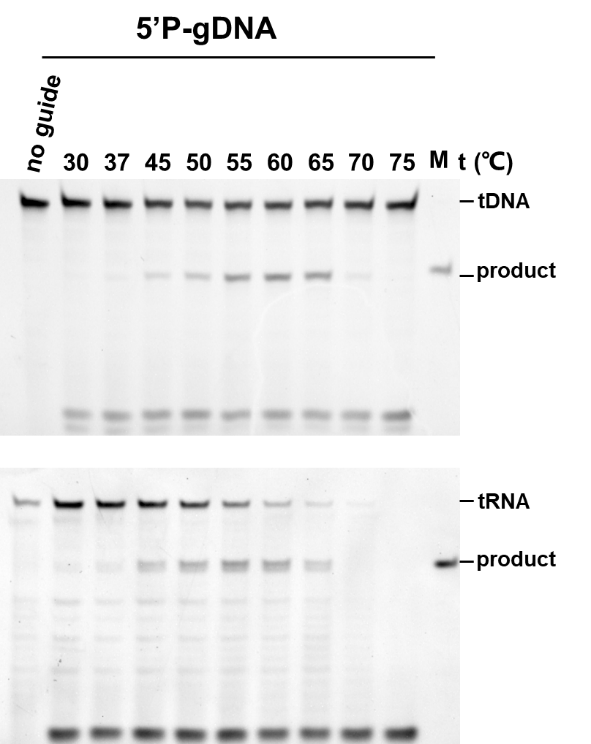

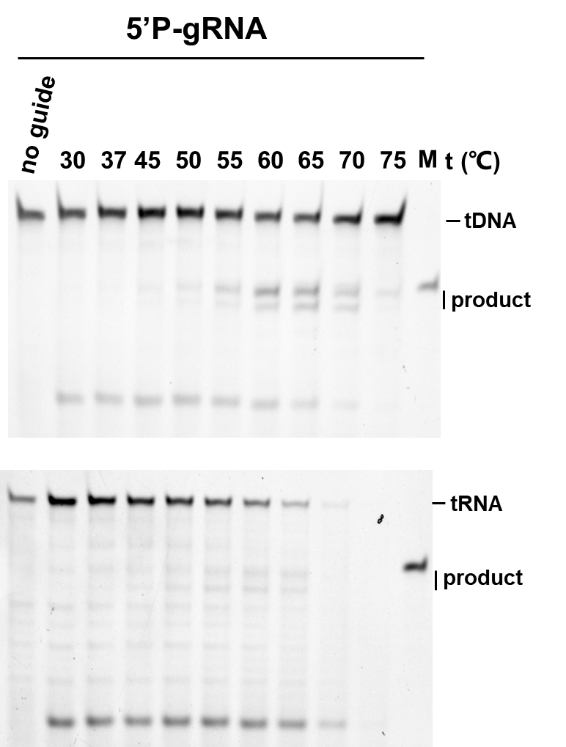

Supplementary Figure S3. Representative denaturing PAGEs (one of three independent experiments) showing MhAgo activity in different temperatures. (A) Representative denaturing PAGEs showing effects of temperatures on ssDNA and RNA cleavage activity mediated by 5’OH-gDNA. (B) Representative denaturing PAGEs showing effects of temperatures on ssDNA and RNA cleavage activity mediated by 5’OH-gRNA. (C) Effects of temperatures on MhAgo activity guided by 5’P-DNA. (D) Effects of temperatures on MhAgo activity guided by 5’P-RNA. The cleavage experiments were performed at the 4:2:1 MhAgo:guide:target molar ratio in reaction buffer containing 5 mM Mn^2+^ ions for 30 min. The no Ago reaction was carried out at 60 °C. The ssDNA targets are completely complementary to corresponding guides the 5’-end nucleotides of which are C. The RNA targets are completely complementary to corresponding guides the 5’-end nucleotides of which are T or U. M indicates synthesized 34-nt ssDNA or RNA product.

**A**

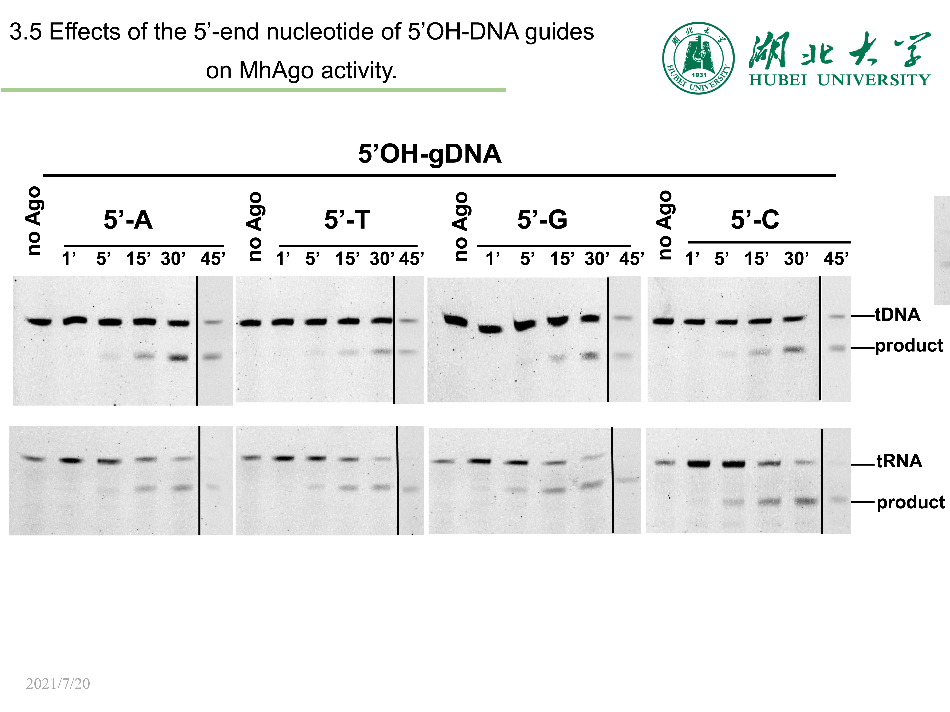

**B**

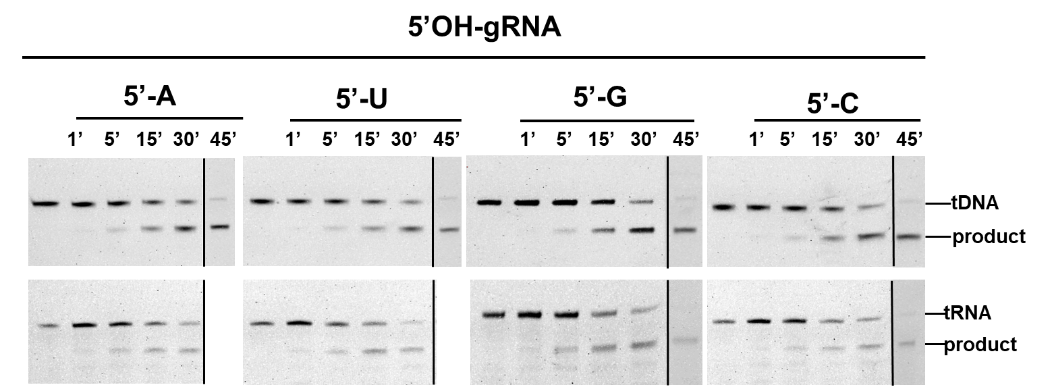

**C**

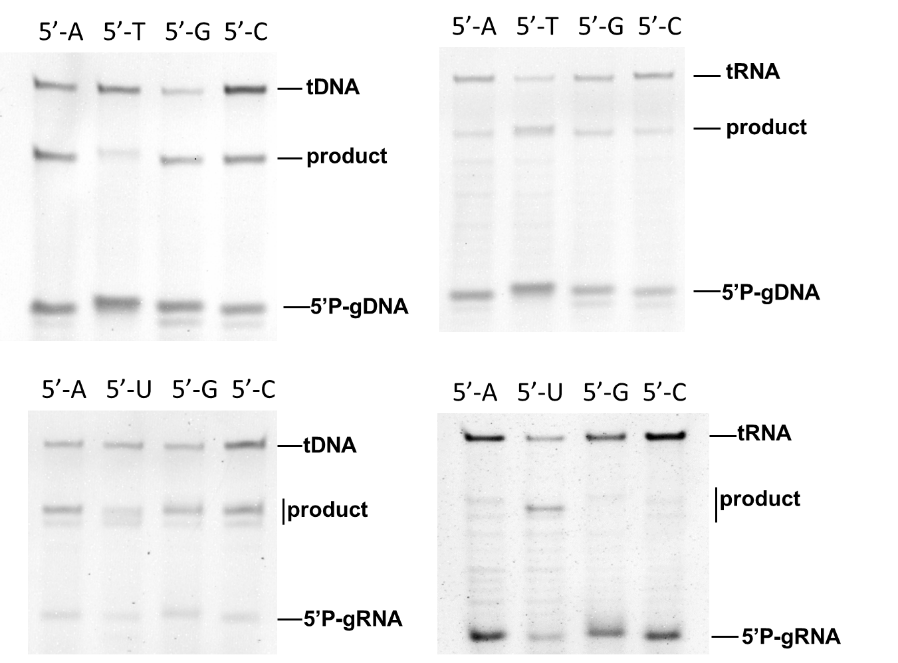

Supplementary Figure S4. Representative denaturing PAGEs (one of three independent experiments) showing effects of the 5’-end nucleotides of guides on MhAgo cleavage activity. (A) Representative denaturing PAGEs showing effects of the 5’-end nucleotides of 5’OH-gDNA on MhAgo cleavage activity. (B) Representative denaturing PAGEs showing effects of the 5’-end nucleotides of 5’OH-gRNA on MhAgo cleavage activity. (C) Representative denaturing PAGEs showing effects of the 5’-end nucleotides of 5’P-guides on MhAgo cleavage activity.

**A**	**B**

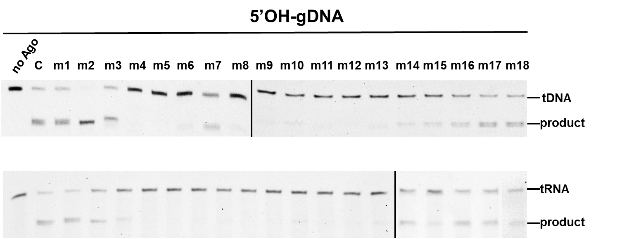

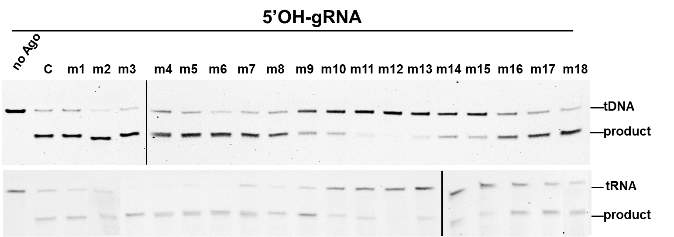

**C**	**D**

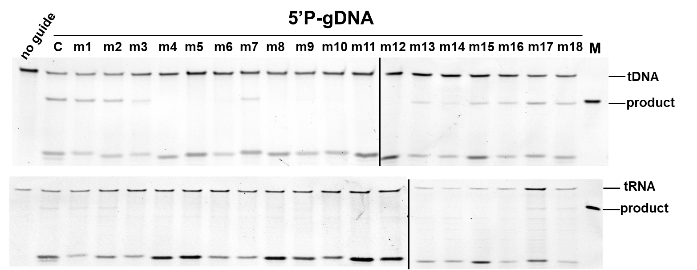

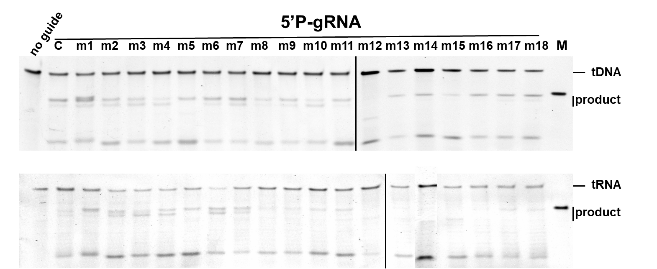

Supplementary Figure S5. Representative denaturing PAGEs (one of three independent experiments) showing effects of mismatches in the guide-target duplexes on the slicing activity of MhAgo. (A) Representative denaturing PAGEs showing effects of guide-target mismatches on MhAgo activity mediated by 5’OH-gDNA. (B) Representative denaturing PAGEs showing effects of guide-target mismatches on MhAgo activity mediated by 5’OH-gRNA. (C) Representative denaturing PAGEs showing effects of guide-target mismatches on MhAgo activity mediated by 5’P-gDNA. (D) Representative denaturing PAGEs showing effects of guide-target mismatches on MhAgo activity mediated by 5’P-gRNA. The experiments were performed at the 4:2:1 MhAgo:guide:target molar ratio in reaction buffer containing 5 mM Mn^2+^ ions at 60 °C. The assay in 5’OH-gDNA mediated ssDNA cleavage was performed for 1 h and the assay in other cleavage patterns was performed for 30 min. The ssDNA and RNA targets are completely complementary to corresponding guides the 5’-end nucleotides of which are C.

**A**	**B**

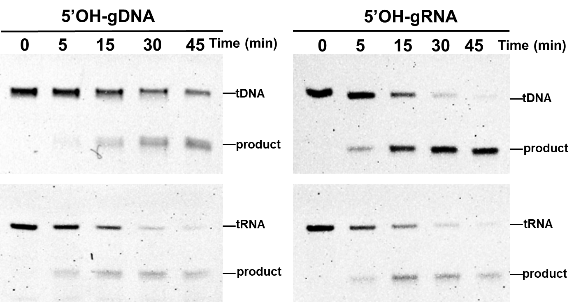

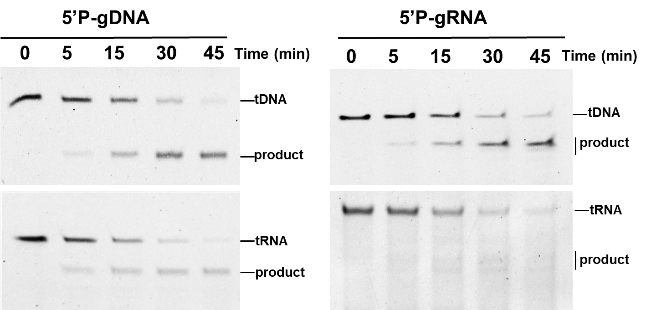

Supplementary Figure S6. Representative denaturing PAGEs showing the cleavage of ssDNA or RNA targets using different guides in a time course experiment. (A) Representative denaturing PAGEs showing the cleavage of ssDNA and RNA targets guided by 5’OH-DNA and 5’OH-RNA in a time course experiment. (B) Representative denaturing PAGEs showing the cleavage of ssDNA and RNA targets guided by 5’P-DNA and 5’P-RNA in a time course experiment.

**A**	**B**

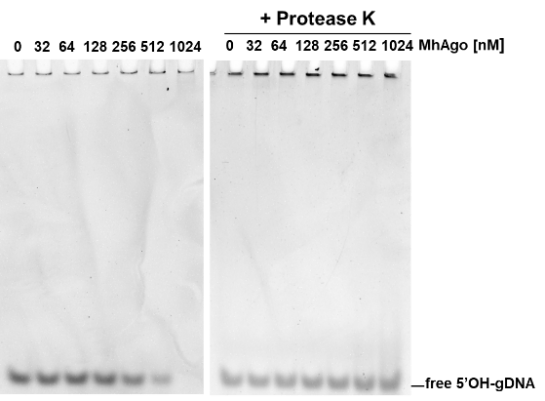

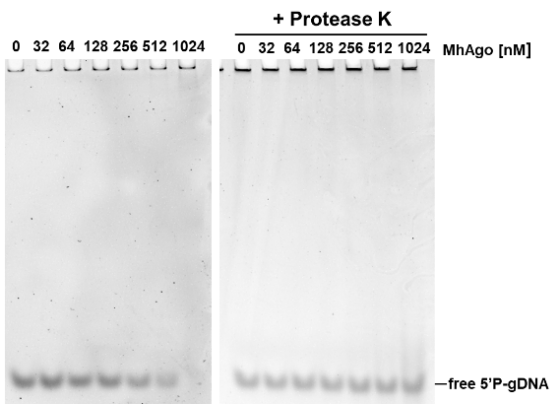

**C**	**D**

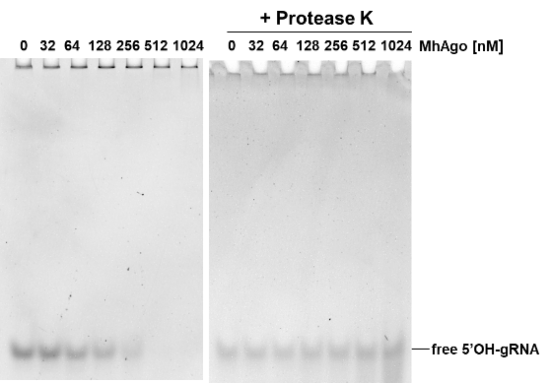

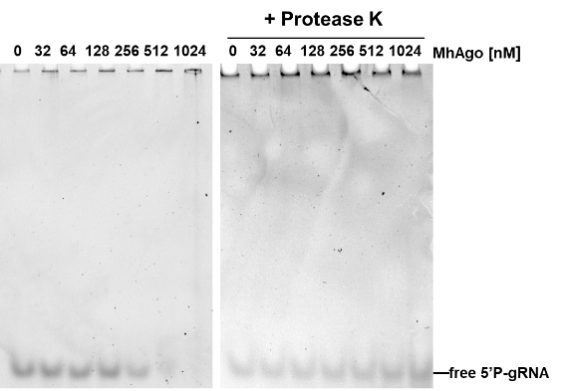

Supplementary Figure S7. Electrophoresis mobility shift assay (EMSA) of the binding of the MhAgo to guides. (A) Representative non-denaturing PAGE of binding reactions with various ratios between MhAgo and 5’OH-gDNA. (B) Representative non-denaturing PAGE of binding reactions with various ratios between MhAgo and 5’P-gDNA. (C) Representative non-denaturing PAGE of binding reactions with various ratios between MhAgo and 5’OH-gRNA. (D) Representative non-denaturing PAGE of binding reactions with various ratios between MhAgo and 5’P-gRNA. All experiments were performed in a binding buffer containing 5 mM Mn^2+^ for 30 min at 55 °C. The samples of binding reactions were treated with protease K for 15 min at 55 °C.

**A**

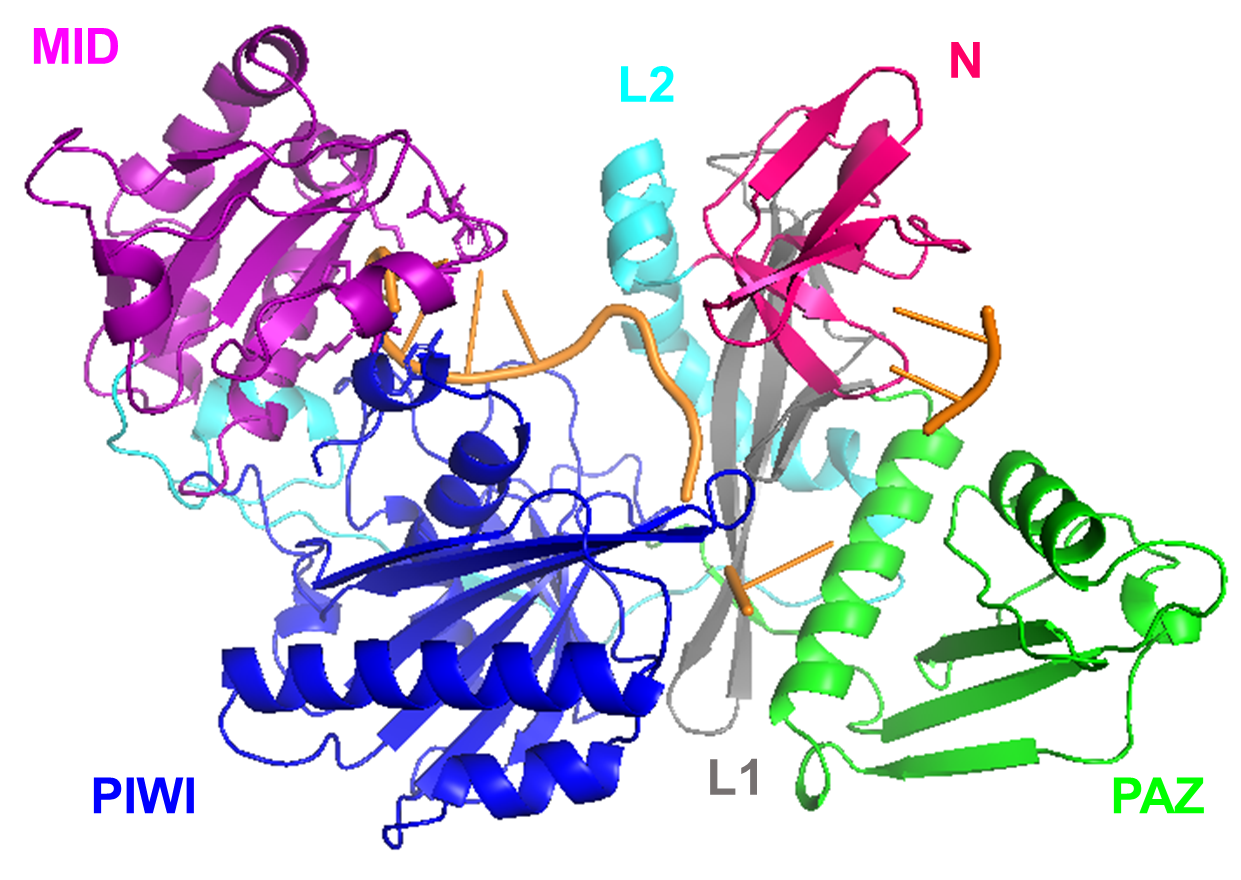
**B**

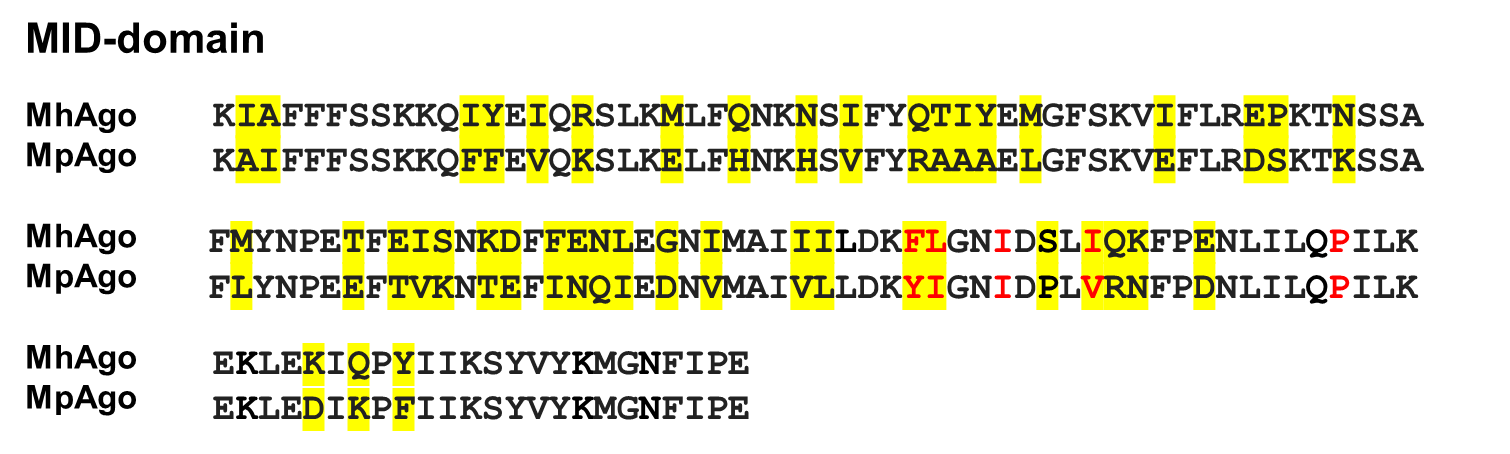

**C**

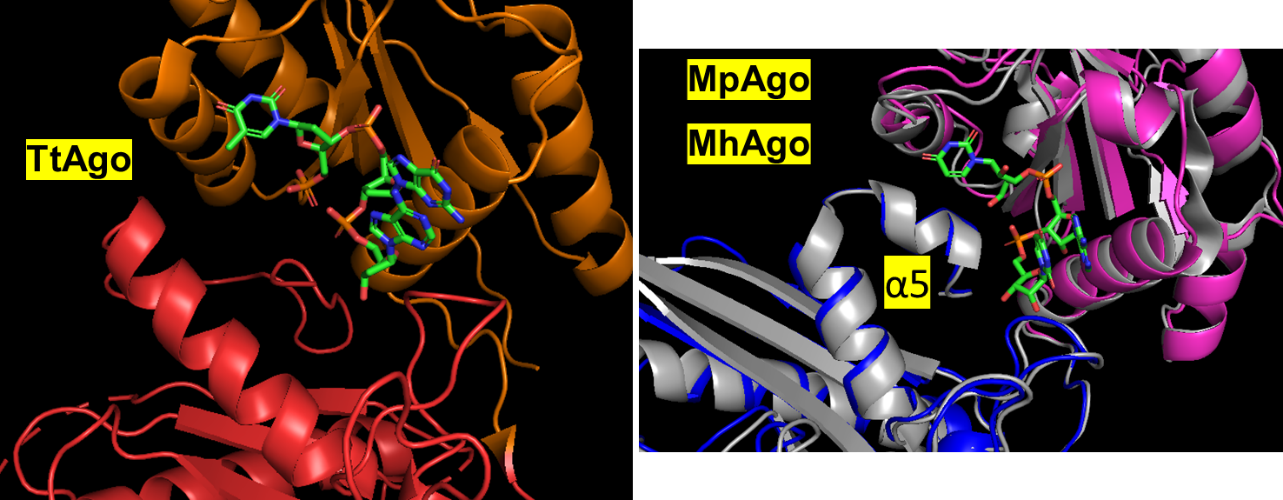

**D**

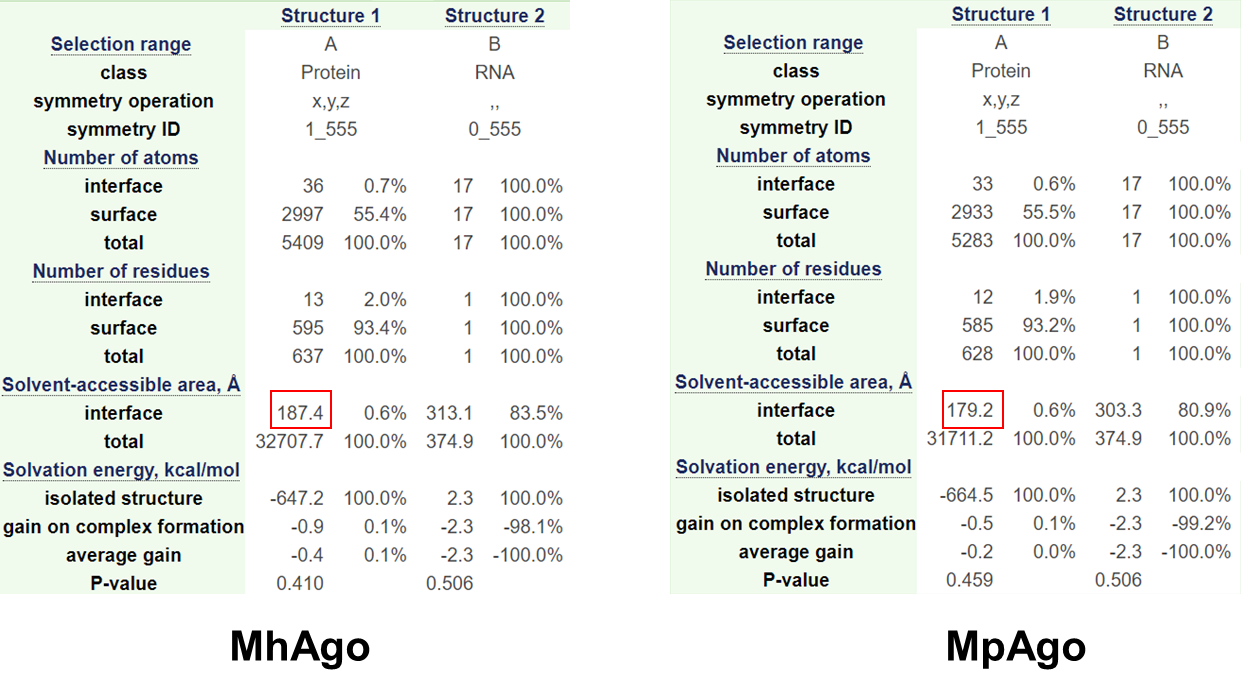

Supplementary Figure S8. The simulated structure analysis of MhAgo. (A) A simulated structure of MhAgo in complex with gRNA (orange) with N (magenta), Linker L1 (gray), PAZ (green), Linker L2 (cyan), MID (purple), and PIWI (blue) domains in a cartoon representation, which is obtained by using the structure of MpAgo in complex with gRNA as a template. (B) Alignment of Mid-domains MhAgo with MpAgo and the different amino acid residues highlighted in yellow. The amino acid residues interacting with 5’-end nucleotide are colored red. (C) In MpAgo (PDB ID: 5I4A, grey) and MhAgo (MID, purple; PIWI, blue), an additional helix (α5) of PIWI domain compresses the MID domain compared with the PIWI domain in TtAgo (PDB ID: 3DLH, MID, orange; PIWI, red). (D) The predicted binding pocket of MhAgo/MpAgo for the 5’-end nucleotide of guide. The size of the binding pocket is indicated by a red box. The size of the binding pocket was calculated with the website https://www.ebi.ac.uk/pdbe/pisa/pistart.html.

**A	B**

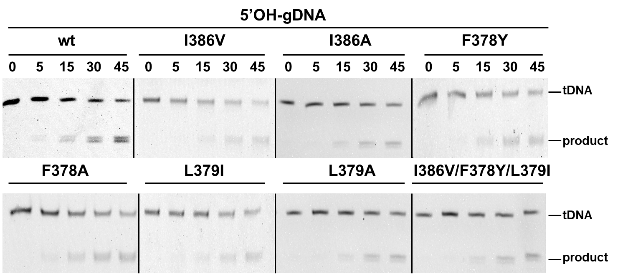

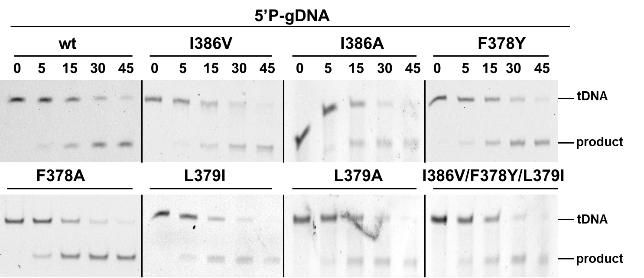

**C	D**

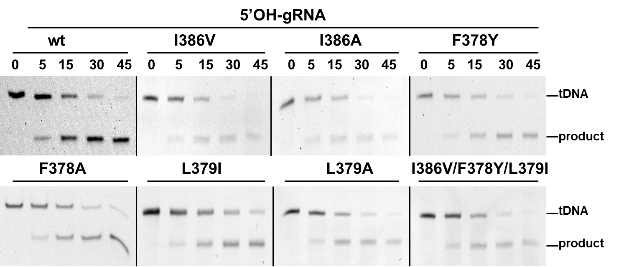

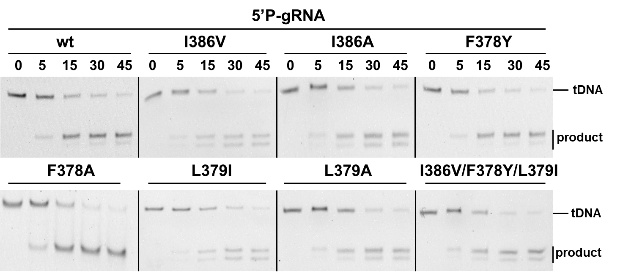

Supplementary Figure S9. The cleavage activity assays of Wild-type MhAgo and its mutants in a time course experiment. (A) Wild-type MhAgo and its mutants are tested for 5’OH-gDNA mediated cleavage activity in a time course experiment. (B) Wild-type MhAgo and its mutants are tested for 5’P-gDNA mediated cleavage activity in a time course experiment. (C) Wild-type MhAgo and its mutants are tested for 5’OH-gRNA mediated cleavage activity in a time course experiment. (D) Wild-type MhAgo and its mutants are tested for 5’P-gRNA mediated cleavage activity in a time course experiment.

**A	B**

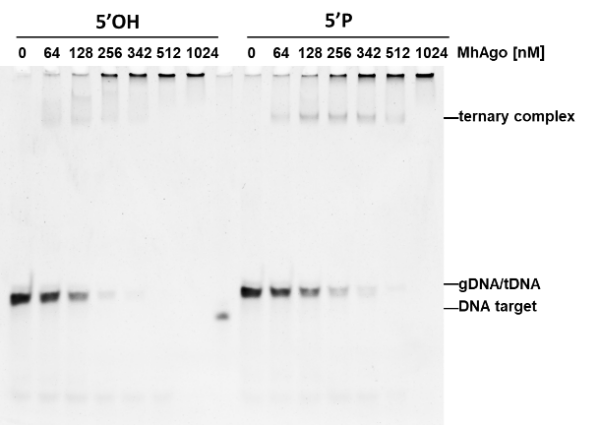

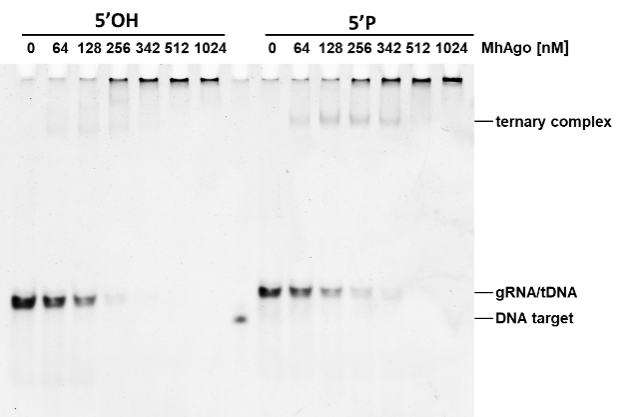

**C	D**

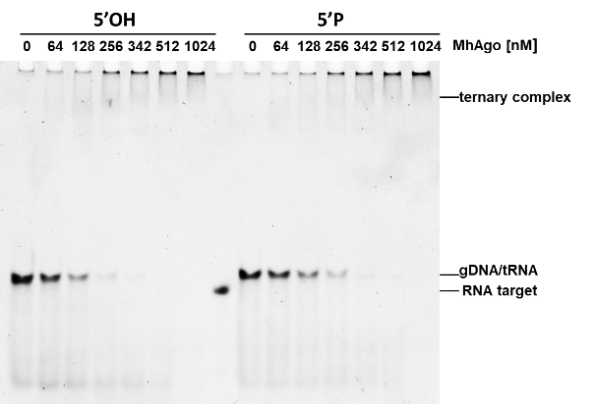

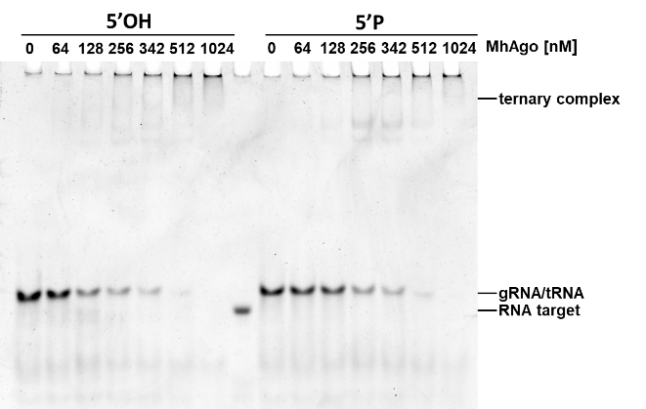

Supplementary Figure S10. Electrophoresis mobility shift assay (EMSA) of the binding of MhAgo to guide: target duplexes. (A) Representative non-denaturing PAGE of binding reactions with various ratios between MhAgo and gDNA:tDNA duplexes. (B) Representative non-denaturing PAGE of binding reactions with various ratios between MhAgo and gRNA:tDNA duplexes. (C) Representative non-denaturing PAGE of binding reactions with various ratios between MhAgo and gDNA:tRNA duplexes. (D) Representative non-denaturing PAGE of binding reactions with various ratios between MhAgo and gRNA:tRNA duplexes.

**Table S1 The cleavage patterns of reported Agos.**

| Name | Cleavage Patterns | | | | | | | | Reaction  Temperature |
| --- | --- | --- | --- | --- | --- | --- | --- | --- | --- |
|  | tDNA | | | | tRNA | | | |  |
|  | **5’OH-gDNA** | **5’P-**  **gDNA** | **5’OH-gRNA** | **5’P-**  **gRNA** | **5’OH-gDNA** | **5’P-**  **gDNA** | **5’OH-gRNA** | **5’P-**  **gRNA** |  |
| **hAgo2** |  | - |  | - |  | + | + | + | 37 °C |
| **KpAgo** |  | - |  | - |  | + |  | + | 30 °C |
| **RsAgo** |  |  |  | - |  |  |  |  | 37 °C |
| **PfAgo** | - | + |  | - |  | - |  | - | 95 °C |
| **MjAgo** | - | + |  | - |  | - |  | - | 95 °C |
| **MpAgo** | - | - | + | + | - | - | + | + | 60 °C |
| **LrAgo** | + | + |  | - |  | - |  | - | 37 °C |
| **CbAgo** | + | + |  | - |  | - |  | - | 37 °C |
| **CbcAgo** |  | + |  |  |  |  |  |  | 37 °C |
| **CpAgo** | + | + | - | - | + | + | - | - | 37 °C |
| **IbAgo** | + | + | - | - | - | - | - | - | 37 °C |
| **KmAgo** | + | + | - | + | + | + | - | + | 55 °C |
| **TtAgo** | - | + |  | - |  | + |  | - | 75 °C |
| **SeAgo** | + | + |  | - |  | - |  | - | 37 °C |
| **TpAgo** |  |  | + | + |  |  |  |  | 60 °C |
| **AaAgo** |  |  |  |  |  | + |  | + | 55 °C |
| **NgAgo** | - | - | - | - | + | + | - | - | 37 °C |
| **FpAgo** | - | + | - | - | - | - | - | - | 95 °C |

The related references have been mentioned in the text.

**Table S2. The nucleic acid sequences used in cleavage and binding experiments.**

| Oligonucleotide name | Sequence (5'-3') | Description |
| --- | --- | --- |
| FAM-A-tDNA | FAM- AAACGACGGCCAGTGCCAAGCTTACTATACAACCTACTACCTCTT | 5’FAM labeled A-tDNA |
| FAM-T-tDNA | FAM- AAACGACGGCCAGTGCCAAGCTTACTATACAACCTACTACCTCAT | 5’FAM labeled T-tDNA |
| FAM-G-tDNA | FAM- AAACGACGGCCAGTGCCAAGCTTACTATACAACCTACTACCTCCT | 5’FAM labeled G-tDNA |
| FAM-C-tDNA | FAM-  AAACGACGGCCAGTGCCAAGCTTACTATACAACCTACTACCTCGT | 5’FAM labeled C-tDNA |
| FAM-A-tRNA | FAM- AAACGACGGCCAGUGCCAAGCUUACUAUACAACCUACUACCUCUU | 5’FAM labeled A-tRNA |
| FAM-T-tRNA | FAM- AAACGACGGCCAGUGCCAAGCUUACUAUACAACCUACUACCUCAU | 5’FAM labeled T-tRNA |
| FAM-G-tRNA | FAM- AAACGACGGCCAGUGCCAAGCUUACUAUACAACCUACUACCUCCU | 5’FAM labeled G-tRNA |
| FAM-C-tRNA | FAM- AAACGACGGCCAGUGCCAAGCUUACUAUACAACCUACUACCUCGU | 5’FAM labeled C-tRNA |
| A-gDNA | AGAGGTAGTAGGTTGTAT | DNA guide forms 5'-A pair with A-tDNA/A-tRNA |
| T-gDNA | TGAGGTAGTAGGTTGTAT | DNA guide forms 5'-T pair with T-tDNA/T-tRNA |
| G-gDNA | GGAGGTAGTAGGTTGTAT | DNA guide forms 5'-G pair with G-tDNA/G-tRNA |
| C-gDNA | CGAGGTAGTAGGTTGTAT | DNA guide forms 5'-C pair with C-tDNA/C-tRNA |
| A-gRNA | AGAGGUAGUAGGUUGUAU | RNA guide forms 5'-A pair with A-tDNA/A-tRNA |
| U-gRNA | UGAGGUAGUAGGUUGUAU | RNA guide forms 5'-T pair with T-tDNA/T-tRNA |
| G-gRNA | GGAGGUAGUAGGUUGUAU | RNA guide forms 5'-G pair with G-tDNA/G-tRNA |
| C-gRNA | CGAGGUAGUAGGUUGUAU | RNA guide forms 5'-C pair with C-tDNA/C-tRNA |
| M1 | FAM- AAACGACGGCCAGTGCCAAGCTTACTATACAACC | 5’ FAM labeled 34 nt DNA |
| M2 | FAM-  AAACGACGGCCAGUGCCAAGCUU ACUAUACAACC | 5’ FAM labeled 34 nt RNA |
| 33nt DNA product | AAACGACGGCCAGTGCCAAGCTTACTATACAAC | 33 nt DNA marker |
| 34nt DNA product | AAACGACGGCCAGTGCCAAGCTTACTATACAACC | 34 nt DNA marker |
| 35nt DNA product | AAACGACGGCCAGTGCCAAGCTTACTATACAACCT | 35 nt DNA marker |
| 33nt RNA product | AAACGACGGCCAGUGCCAAGCUUACUAUACAAC | 33 nt RNA marker |
| 34nt RNA product | AAACGACGGCCAGUGCCAAGCUUACUAUACAACC | 34 nt RNA marker |
| 35nt RNA product | AAACGACGGCCAGUGCCAAGCUUACUAUACAACCU | 35 nt RNA marker |
| C-tDNA | AAACGACGGCCAGTGCCAAGCTTACTATACAACCTACTACCTCGT | 45 nt DNA target for C-gDNA/C-gRNA |
| C-tRNA | AAACGACGGCCAGUGCCAAGCUUACUAUACAACCUACUACCUCGU | 45 nt RNA target for A-gDNA/A-gRNA |
| gDNA_mm1 | GGAGGTAGTAGGTTGTAT | DNA guide forms mismatched pair in position 1 with C-tDNA/C-tRNA |
| gDNA_mm2 | CCAGGTAGTAGGTTGTAT | DNA guide forms mismatched pair in position 2 with C-tDNA/C-tRNA |
| gDNA_mm3 | CGTGGTAGTAGGTTGTAT | DNA guide forms mismatched pair in position 3 with C-tDNA/C-tRNA |
| gDNA_mm4 | CGACGTAGTAGGTTGTAT | DNA guide forms mismatched pair in position 4 with C-tDNA/C-tRNA |
| gDNA_mm5 | CGAGCTAGTAGGTTGTAT | DNA guide forms mismatched pair in position 5 with C-tDNA/C-tRNA |
| gDNA_mm6 | CGAGGAAGTAGGTTGTAT | DNA guide forms mismatched pair in position 6 with C-tDNA/C-tRNA |
| gDNA_mm7 | CGAGGTTGTAGGTTGTAT | DNA guide forms mismatched pair in position 7 with C-tDNA/C-tRNA |
| gDNA_mm8 | CGAGGTACTAGGTTGTAT | DNA guide forms mismatched pair in position 8 with C-tDNA/C-tRNA |
| gDNA_mm9 | CGAGGTAGAAGGTTGTAT | DNA guide forms mismatched pair in position 9 with C-tDNA/C-tRNA |
| gDNA_mm10 | CGAGGTAGTTGGTTGTAT | DNA guide forms mismatched pair in position 10 with C-tDNA/C-tRNA |
| gDNA_mm11 | CGAGGTAGTACGTTGTAT | DNA guide forms mismatched pair in position 11 with C-tDNA/C-tRNA |
| gDNA_mm12 | CGAGGTAGTAGCTTGTAT | DNA guide forms mismatched pair in position 12 with C-tDNA/C-tRNA |
| gDNA_mm13 | CGAGGTAGTAGGATGTAT | DNA guide forms mismatched pair in position 13 with C-tDNA/C-tRNA |
| gDNA_mm14 | CGAGGTAGTAGGTAGTAT | DNA guide forms mismatched pair in position 14 with C-tDNA/C-tRNA |
| gDNA_mm15 | CGAGGTAGTAGGTTCTAT | DNA guide forms mismatched pair in position 15 with C-tDNA/C-tRNA |
| gDNA_mm16 | CGAGGTAGTAGGTTGAAT | DNA guide forms mismatched pair in position 16 with C-tDNA/C-tRNA |
| gDNA_mm17 | CGAGGTAGTAGGTTGTTT | DNA guide forms mismatched pair in position 17 with C-tDNA/C-tRNA |
| gDNA_mm18 | CGAGGTAGTAGGTTGTAA | DNA guide forms mismatched pair in position 18 with C-tDNA/C-tRNA |
| gRNA_mm1 | GGAGGUAGUAGGUUGUAU | RNA guide forms mismatched pair in position 1 with C-tDNA/C-tRNA |
| gRNA_mm2 | CCAGGUAGUAGGUUGUAU | RNA guide forms mismatched pair in position 2 with C-tDNA/C-tRNA |
| gRNA_mm3 | CGUGGUAGUAGGUUGUAU | RNA guide forms mismatched pair in position 3 with C-tDNA/C-tRNA |
| gRNA_mm4 | CGACGUAGUAGGUUGUAU | RNA guide forms mismatched pair in position 4 with C-tDNA/C-tRNA |
| gRNA_mm5 | CGAGCUAGUAGGUUGUAU | RNA guide forms mismatched pair in position 5 with C-tDNA/C-tRNA |
| gRNA_mm6 | CGAGGAAGUAGGUUGUAU | RNA guide forms mismatched pair in position 6 with C-tDNA/C-tRNA |
| gRNA_mm7 | CGAGGUUGUAGGUUGUAU | RNA guide forms mismatched pair in position 7 with C-tDNA/C-tRNA |
| gRNA_mm8 | CGAGGUACUAGGUUGUAU | RNA guide forms mismatched pair in position 8 with C-tDNA/C-tRNA |
| gRNA_mm9 | CGAGGUAGAAGGUUGUAU | RNA guide forms mismatched pair in position 9 with C-tDNA/C-tRNA |
| gRNA_mm10 | CGAGGUAGUUGGUUGUAU | RNA guide forms mismatched pair in position 10 with C-tDNA/C-tRNA |
| gRNA_mm11 | CGAGGUAGUACGUUGUAU | RNA guide forms mismatched pair in position 11 with C-tDNA/C-tRNA |
| gRNA_mm12 | CGAGGUAGUAGCUUGUAU | RNA guide forms mismatched pair in position 12 with C-tDNA/C-tRNA |
| gRNA_mm13 | CGAGGUAGUAGGAUGUAU | RNA guide forms mismatched pair in position 13 with C-tDNA/C-tRNA |
| gRNA_mm14 | CGAGGUAGUAGGUAGUAU | RNA guide forms mismatched pair in position 14 with C-tDNA/C-tRNA |
| gRNA_mm15 | CGAGGUAGUAGGUUCUAU | RNA guide forms mismatched pair in position 15 with C-tDNA/C-tRNA |
| gRNA_mm16 | CGAGGUAGUAGGUUGAAU | RNA guide forms mismatched pair in position 16 with C-tDNA/C-tRNA |
| gRNA_mm17 | CGAGGUAGUAGGUUGUUU | RNA guide forms mismatched pair in position 17 with C-tDNA/C-tRNA |
| gRNA_mm18 | CGAGGUAGUAGGUUGUAA | RNA guide forms mismatched pair in position 18 with C-tDNA/C-tRNA |
| 12nt C-gDNA | CGAGGTAGTAGG | 12 nt DNA guide pair with C-tDNA/C-tRNA |
| 13nt C-gDNA | CGAGGTAGTAGGT | 13 nt DNA guide pair with C-tDNA/C-tRNA |
| 14nt C-gDNA | CGAGGTAGTAGGTT | 14 nt DNA guide pair with C-tDNA/C-tRNA |
| 15nt C-gDNA | CGAGGTAGTAGGTTG | 15 nt DNA guide pair with C-tDNA/C-tRNA |
| 16nt C-gDNA | CGAGGTAGTAGGTTGT | 16 nt DNA guide pair with C-tDNA/C-tRNA |
| 17nt C-gDNA | CGAGGTAGTAGGTTGTA | 17 nt DNA guide pair with C-tDNA/C-tRNA |
| 19nt C-gDNA | CGAGGTAGTAGGTTGTATA | 19 nt DNA guide pair with C-tDNA/C-tRNA |
| 20nt C-gDNA | CGAGGTAGTAGGTTGTATAG | 20 nt DNA guide pair with C-tDNA/C-tRNA |
| 21nt C-gDNA | CGAGGTAGTAGGTTGTATAGT | 21 nt DNA guide pair with C-tDNA/C-tRNA |
| 25nt C-gDNA | CGAGGTAGTAGGTTGTATAGTAAGC | 25 nt DNA guide pair with C-tDNA/C-tRNA |
| 30nt C-gDNA | CGAGGTAGTAGGTTGTATAGTAAGCTTGGC | 30 nt DNA guide pair with C-tDNA/C-tRNA |
| 12nt C-gRNA | CGAGGUAGUAGG | 12 nt RNA guide pair with C-tDNA/C-tRNA |
| 13nt C-gRNA | CGAGGUAGUAGGU | 13 nt RNA guide pair with C-tDNA/C-tRNA |
| 14nt C-gRNA | CGAGGUAGUAGGUU | 14 nt RNA guide pair with C-tDNA/C-tRNA |
| 15nt C-gRNA | CGAGGUAGUAGGUUG | 15 nt RNA guide pair with C-tDNA/C-tRNA |
| 16nt C-gRNA | CGAGGUAGUAGGUUGU | 16 nt RNA guide pair with C-tDNA/C-tRNA |
| 17nt C-gRNA | CGAGGUAGUAGGUGUA | 17 nt RNA guide pair with C-tDNA/C-tRNA |
| 19nt C-gRNA | CGAGGUAGUAGGUUGUAUA | 19 nt RNA guide pair with C-tDNA/C-tRNA |
| 20nt C-gRNA | CGAGGUAGUAGGUUGUAUAG | 20 nt RNA guide pair with C-tDNA/C-tRNA |
| 21nt C-gRNA | CGAGGUAGUAGGUUGUAUAGU | 21 nt RNA guide pair with C-tDNA/C-tRNA |
| 25nt C-gRNA | CGAGGUAGUAGGUUGUAUAGUAAGC | 25 nt RNA guide pair with C-tDNA/C-tRNA |
| 30nt C-gRNA | CGAGGUAGUAGGUUGUAUAGUAAGCUUGGC | 30 nt RNA guide pair with C-tDNA/C-tRNA |
